# Morphological and microbial diversity of hydromagnesite microbialites in Lake Salda, A Ma8rs Analog Alkaline Lake

**DOI:** 10.1101/2023.09.06.556485

**Authors:** Yagmur Gunes, Fatih Sekerci, Burak Avcı, Thijs J. G. Ettema, Nurgul Balci

**Affiliations:** Department of Geological Engineering, Geomicrobiology & Biogeochemistry Laboratory, Istanbul Technical University, Istanbul, Turkey; Laboratory of Microbiology, Wageningen University and Research, Wageningen, The Netherland; Department of Geology, University of Georgia, Athens, GA, USA

**Keywords:** Microbialites, Hydromagnesite, Terrestrial analog, Lake Salda, Prokaryotic Diversity, *Exiguobacterium*

## Abstract

Lake Salda, recognized as a terrestrial analog for the paleolake in Jezero Crater on Mars, hosts modern, sub/fossil and fossil hydromagnesite microbialites. A comprehensive study was conducted to reveal the distribution, and morphological-mineralogical, microbial diversity of the microbialites that are currently growing in the shallow (<1 m) and deeper waters (up to 15 m) of the lake. Six major microbialite forming zones were identified (Zone I-VI) comprising previously unknown morphotypes of the microbialites. These newly identified morphologies include; linked columns and microbial pavements, exhibiting distinct surface textures such as bulbous, mini columnar, knobby, cerebroidal and smooth. The shallow microbialites showed well-preserved radially growing stromatolitic layers in cm-scale, producing cauliflower structures while the deeper samples exhibit mm scale layering within the mini columnar structures. 16S rRNA amplicon data reveal that abundance of major bacterial taxa of the thrombolitic microbialites from deeper regions are distinct than those of the shallow-growing microbialites with stromatolitic structures. Cyanobacteria were found to be generally less abundant (0.01% to 2%) in the thrombolitic microbialites compared to the stromatolitic microbialites with smooth surface. The Anaerolineae class of the phylum Chloroflexi was prominently present in the stromatolitic microbialites with smooth surface. Notably, the lake exhibited a high abundance of the genus *Exiguobacterium*, particularly in the deep thrombolitic microbialites (e.g., 93%). Microscopy analysis of the deep microbial mats showed the presence of abundant nano and microcrystalline platy/flakey hydrated Mg carbonate crystals that co- existed with mineralized filaments and spherulitic aggregates within abundant exopolymeric substances (EPSs). Moreover, palygorskite mineral was exclusively identified within the deep microbialites. Co-existence of aragonite and hydromagnesite minerals in particular within the deep microbialites that grow under non-evaporative conditions and the abundant presence of entombed biomass (e.g., filamentous) collectively suggest the potential preservation of biosignatures within the hydrated Mg carbonate build-ups in Lake Salda.

## INTRODUCTION

Microbialites are carbonate structures that are built or influenced by benthic microorganisms. They have been a prominent feature throughout Earth’s history in a wide range of geological environments from Precambrian carbonate platforms (Hofmann *et al.,* 1999; Allwood *et al.,* 2004; Murphy and Sumner, 2008) to contemporary Mg-carbonate build-ups (Frantz *et al.,* 2015; Chagas *et al.,* 2016). These attached, lithified, organo-sedimentary structures represent the oldest life form on Earth, with preserving records dating back over 3 billion years (Awramik & Margulis, 1974; Van Kranendonk *et al.,* 2008). These oldest putative macrofossils offer valuable insights into the geochemical, environmental, and biological records of early Earth. Furthermore, the study of carbonates in microbialites has significant implications for the detection and understanding of similar formations on Mars, providing clues about the potential existence of past life on the Red Planet.

Orbital spectroscopy (Mars Reconnaissance Orbiter’s Compact Reconnaissance Imaging Spectrometer, CRISM) has identified olivine, pyroxene, and Mg-carbonates, in the marginal deposits of the paleolake in Jezero crater, Mars (Horgan *et al*., 2020, Tarnas *et al*., 2021). In particular, strong spectral signals resembling hydrated Mg carbonates - hydromagnesite [Mg_5_(CO_3_)4(OH)_2_.4H_2_O] - have been received from the marginal carbonates in the delta and along the littoral zone of the paleolake basin (Horgan *et al*., 2020). These carbonate-bearing deposits in the Jezero Crater could indicate detrital sedimentation in the delta and some near-shore precipitation of hydromagnesite along the crater’s margin. The identified mineralogy and depositional features of the carbonates in the Jezero Crater make the marginal carbonates, in particular within the littoral zone of the paleolake, the prime target for the search for biosignatures.

Carbonates that precipitate with often biological mediation within lacustrine systems, as extensively documented on Earth, possess high potential for preserving morphological, organic, and isotopic biosignatures (Braissant *et al*., 2007; Theisen *et al*., 2015; Chagas *et al*., 2016 and references therein.). Thus, the Mg-carbonate bearing deposits in Jezero may also have high biosignature preservation potential as those on Earth. However, our ability to identify and characterize facies with high biosignature preservation potential in a microbially-dominated, mafic, and alkaline lacustrine system hosting hydrated Mg carbonates that have been hypothesized for the Jezero paleolake is currently limited. Such systems are rare on Earth today as well as in the geological record, posing challenges for understanding and interpreting the potential biosignatures in the Jezero Crater.

Modern terrestrial analogs of lacustrine carbonates on Mars can be found in lacustrine systems within ultramafic terrains. These analogs provide valuable opportunity to expand our understanding of carbonate deposition at Jezero Crater, which is, a crucial step towards to determining paleolake processes and refining sampling strategies to target the most promising areas for biosignatures preservations. One such system is Lake Salda, Turkey (Fig. 1) where fractured ultramafic rocks are exposed to the flow of surface and subsurface fluids, leading to the formation of contemporary hydrated Mg carbonate microbialites along the lake’s perimeter (Balci *et al*., 2020, Garczynski *et al*., 2020).

**Figure 1.**
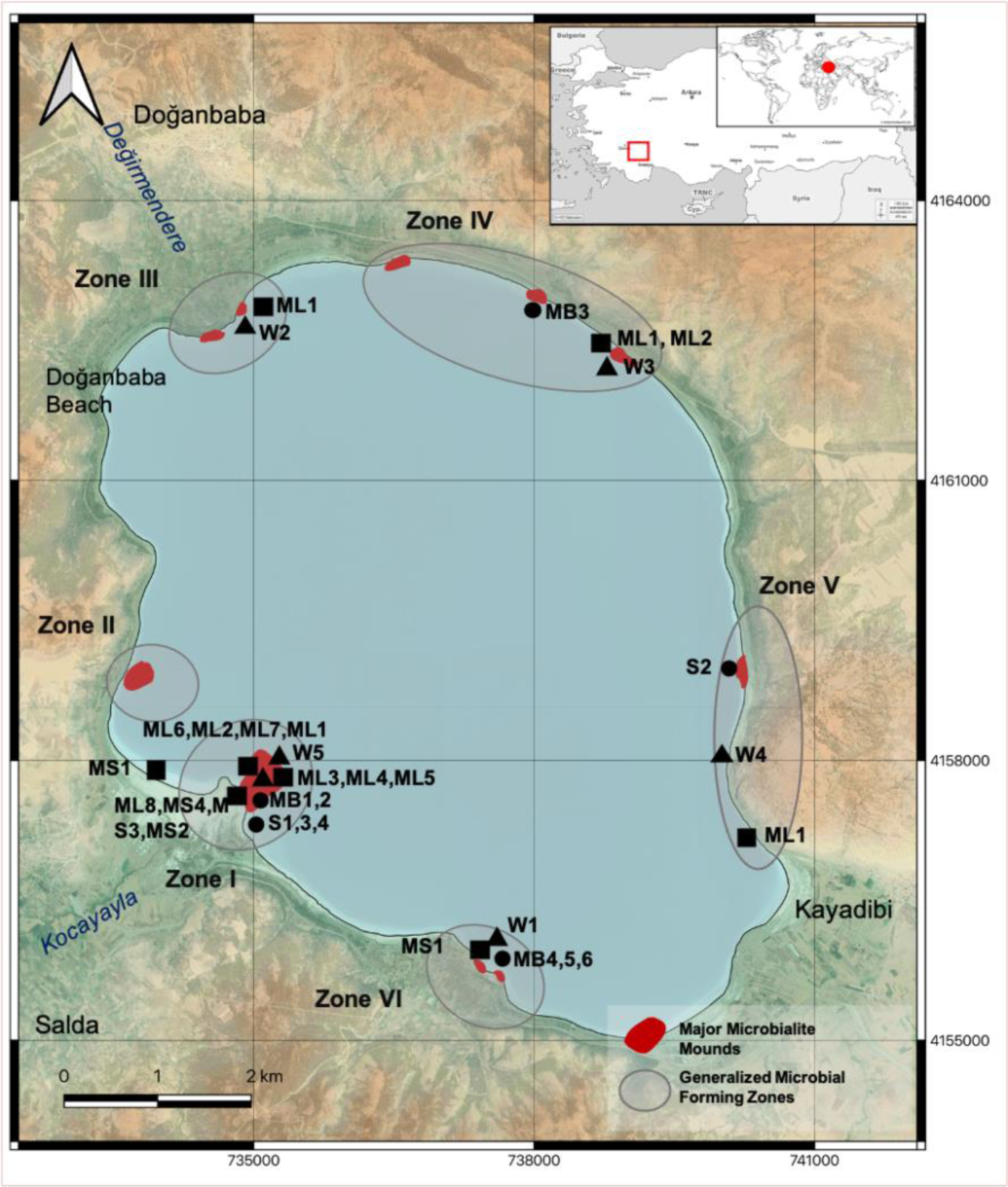
Locations of major microbialite forming zones (I-VI) and the field samplings including water, sediment and microbialites: ▴: Hydrochemistry, ▪: Geochemistry, ●: Prokaryotic diversity.

Lake Salda offers an exceptional setting to investigate the microbially mediated precipitation of hydrated magnesium carbonates in a lacustrine system due to the morphological and microbial diversity of microbialites. Lacustrine environments have various geochemical conditions, including freshwater, evaporative and alkaline, and hypersaline which give rise to microbialites with distinct morphologies (e.g., stromatolites, thrombolites, dendrolite, leiolite), and calcium and magnesium compositions, as well as different textures (laminated, clotted etc.) (Kempe and Kaźmierczak 1990; Arp *et al.,* 2003; Bougeault *et al.,* 2019). The relationship between morphological diversity and microbial community profiles of microbialites, in particular mat- forming communities, has been also reported (Knoll et *al.,* 1999; Lopez-Garcia *et al*., 2005; Dupraz *et al*., 2011). Based on microbial diversity profiles, sedimentary structures, and isotopic compositions in modern microbialites, various microbial carbonate fixation processes, such as cyanobacterial, oxygenic photosynthesis (Altermann *et al.,* 2006; Riding 2006), anoxygenic photosynthesis (Bosak *et al.,* 2007) and sulfur metabolism (Wacey *et al.,* 2011), have been proposed to induce the formation of carbonates. These unique features of microbialites in lacustrine environments make them highly valuable for studying and interpreting the records of their ancient analogs on Earth and potentially on Mars.

Previous studies on Lake Salda have primarily focused on geochemical and limnological aspects, with only a few delving into the role of microbiological processes in the formation of microbialites (Braithwaite and Zedef 1994; 1996; Shirokova *et al*., 2013; Balci *et al*., 2018; 2020) and comparing them with carbonates on Mars (Russel *et al*., 1999; Garczynski *et al.,* 2020). Also, there are only a few studies that have explored the microbial diversity of the actively growing microbialites, which are only restricted to the western part of the lake (Poyraz and Mutlu, 2017; Balci *et al*., 2020; Balci 2023). In a recent study, Balci *et al*. (2020) and Balci (2023) demonstrated the significance of EPS production and degradation in microbialite growth in Lake Salda, proposing a conceptual growth model. Other experiments involving isolated cyanobacteria and lipid biomarker characterization have also suggested that the formation of microbialites in Lake Salda has been primarily regulated by a phototrophic community dominated by cyanobacteria and diatoms (Braithwaite and Zedef 1996; Shirokova *et al*., 2013; Mavromatis *et al*., 2015; Kaiser *et al*., 2016; Balci *et al*., 2018; 2020).

Despite the accumulated knowledge regarding the mineralogical and geochemical characteristics of Lake Salda and its vicinity, there is still a lack of information on the microbialite morphology around the lake and its correlation to the site- specific microbial community profiles. In this study, we present, for the first time, a site-specific prokaryotic diversity of the microbialites and further investigate the potential connection between the microbial diversity and morphology of microbialites as well as their depositional settings around the lake.

Novel findings from terrestrial analogues like Lake Salda greatly contribute to our understanding of the interplay of microbial, chemical, and environmental processes that mediate the precipitation of hydrated magnesium carbonates in a lacustrine system and help us gain insights into the nature of carbonate deposition at Jezero Crater, Mars.

## SITE DESCRIPTION

Lake Salda is located in Burdur, SW Turkey, within an endorheic basin (Fig. 1). The lake’s paleo detrital input from the fluvial systems has resulted in the formation of two major fan delta plains on the north (Doganbaba) and the west sides (Salda) of the lake basin. The modern lake system has limited surface water inflow with precipitation, groundwater and ephemeral streams associated with Kocayayla and Degirmendere rivers, serving as the main sources of water (Davraz *et al*., 2019). The only outlet for water from the lake is through evaporation with an average from the lake surface area estimated at 53.98 ×10^6^ m^3^/year (Varol *et al.,* 2018).

The lake water has generally high Mg/Ca ratios ranging from 800 to 3170 with a maximum recorded value of approximately 7000. The pH of the lake water varies from 8.7 and 10.0. The water chemistry of the lake does not significantly fluctuate seasonally and typically shows a decreasing trend of Mg > Na > Si > Ca > K (Varol *et al*., 2020; Balci *et al*., 2020). The lake is classified as oligotrophic with a max. sunlight penetrating depth of 17 m, thus favoring phototrophs to live in the deeper part of the water column (Kazanci *et al*., 2004).

Ultramafic rocks presented by the Marmaris Peridotite complex covers the lake’s immediate area and most of the basin. Dolomitized Cretaceous limestone underlies the eastern part of the lake. In the SE at Yesilova, the SW at Kocaadalar and the NW at Doganbaba, there are alluvial fan deltas that are comprised of gravel and pebbles of serpentinite and gabbro with extensively weathered surfaces. Lake Salda shores vary from low- energy beaches composed of sand or mud flats bordering deltas to steep and rocky shores. Mg enrichment of the Lake Salda water has been attributed to the washing of the peridotite pebbles around the lake (e.g. harzburgite, dunite). Carbonate terrace deposits are observable at the SE and SW beaches of the lake reaching up to 15 m in height.

## MATERIALS AND METHODS

### Field Observations and Sampling

The data presented in this study were obtained during the sampling campaigns between the years 2019 and 2022. In 2022, the lake experienced relatively low rainfall with only 32 cm recorded for the months of February and March. This resulted in a significant drop in the lake level, allowing the sampling of the freshly exposed microbialites. Additional to the actively growing microbialite samples, dried out/sub fossilized samples were also collected. Detailed morphological examinations of the microbialites, as well as the depositional characteristics of the microbialite-hosting environments, were carried out during the excursions. The mapping of microbialite formations along the lake shore was carried out and six major microbialite formation zones were identified (Zone I-VI). Descriptive classifications of the microbialites in each zone based on their macro and mesoscale morphology. Various samples, including sediment, water, and microbialite were collected at each identified zone (Fig.1). Special attention was given to water sampling, specifically targeting areas in the close vicinity of the actively growing microbialites. In–situ measurements of physicochemical properties such as pH, T°C and total dissolved solids (TDS) were conducted using Hanna HI (v.9812-5). Before each measurement, the pH probe was calibrated against pH buffer solutions of 4.01, 7.01 and 11.01. Analysis of standards showed an accuracy of better than ± 0.04 pH units for the 7.0 pH buffer solutions with a precision better than 98.8 %. The conductivity probe exhibited an average error percentage of ± 9.1 % and ± 4.3 % when the standard solutions 147, 1413 and 12880 µS/cm were used, respectively.

Upon collection, all water samples were immediately filtered using a syringe filter (0.22 μm, Sartorious). The filtered water was then stored in pre-rinsed 250 ml HDPE bottles in cold, dark conditions at a temperature of 4°C. For cation analysis, a 100 ml portion of water sample was acidified with ultrapure 3M nitric acid (Merck) and the remaining water was stored for the analysis of ions concentration. To measure alkalinity, the water samples were completely filled in a 1L bottle and the mouth of the bottle was covered with parafilm to prevent air exposure. To determine the water depth relative to the microbialites, a marked plastic pipe was used at locations close to the sampled microbialites. The length between the top surface of the shoreline sediments and the water surface was measured and recorded as the water depth (Table 1).

**Table 1.** Detailed descriptions of microbialite (ML=16) and sediment (n=5) samples subjected to 16S rRNA analysis.

| Zone | Sample Name | Sample type | Depth |  | Color | Lithification | Environmental Conditions | Description |
| --- | --- | --- | --- | --- | --- | --- | --- | --- |
|  |  |  | Sediment | Water |  |  |  |  |
| Zone I | ML1-I | Microbialite |  | 80 cm | Purple | L | Submerged | Layered carbonate crusts within submerged, actively growing microbialite |
|  | ML2-I | Microbialite |  | 20 cm | White | L | Submerged | Layered carbonate crusts within submerged, actively growing microbialite |
|  | ML3-I | Microbial Mat |  | 5m | Bright yellow | L | Submerged | Actively growing submerged microbialite. Uppermost part of the microbial layer |
|  | ML4-I | Microbialite |  | 10 m | Dark purple | L | Submerged | Actively growing submerged microbialite. Carbonate layer with dark purple pigmentation |
|  | ML5-I | Microbial Mat |  | 15 m | Bright yellow | L | Submerged | Uppermost microbial layer of actively growing submerged microbialite. |
|  | ML6-I | Microbial Mat |  | 15 cm | Dark yellow, pink-ish orange | L | Submerged | Uppermost microbial layer of actively growing submerged microbialite |
|  | ML7-I | Microbial Mat |  | 15 cm | Bright yellow | UL | Submerged | Actively growing submerged microbial layer. Uppermost organic layer with fibrous texture. Lower section only has carbonate clasts |
|  | MS1-I | Sediment |  | 10 cm | Gray to pink-ish | UL | Submerged | Beach deposit with unconsolidated clayey sand and mucilaginous texture |
|  | MS2-I | Sediment |  | 20 cm | Gray to pink-ish | UL | Submerged | Beach deposit with mucilaginous texture |
|  | ML8-I | Microbial Mat |  | Surface | Pale yellow | UL | Splash Zone | Microbial layer covering the sediment deposits |

**Table 1 (continued).** Detailed descriptions of microbialite (ML=16) and sediment (n=5) samples subjected to 16S rRNA analysis.
| Zone | Sample Name | Sample type | Depth |  | Color | Lithification | Environmental Conditions | Description |
| --- | --- | --- | --- | --- | --- | --- | --- | --- |
|  |  |  | Sediment | Water |  |  |  |  |
| Zone I | MS3-I | Sediment | 10 cm |  | Gray to pink-ish | UL | Splash Zone | Beach deposit with unconsolidated clay-sand |
|  | MS4-I | Sediment | 15 cm |  | Dark grey, black | UL | Splash Zone | Dark colored beach deposit with mucilaginous texture |
| Zone III | ML1-III | Microbialite |  | Surface | Dark yellow | L | Splash Zone | mm-thick uppermost crust with mm to cm sized vugs. |
| Zone IV | ML2-IV | Microbial Mat |  | 10 cm | Gray to pink-ish | UL | Submerged | Smooth mucilaginous texture overlaid by muddy sand with mm scale crust on top |
| Zone IV | ML1-IV | Microbial Mat |  | 80 cm | Dark yellow | L | Submerged | Actively growing submerged microbialite. Uppermost microbial layer with fibrous texture. |
| Zone V | ML1-V | Microbial Mat |  | 5 cm | Pale yellow | UL | Submerged | Actively growing submerged microbialite. Uppermost microbial layer with fibrous texture. Carbonate clasts on the lower section. |
| Zone VI | MS1-VI | Sediment |  | 80 cm | Grey | L | Submerged | Gabbroic grains with hydromagnesite clasts |

At each water sampling point, whenever possible, microbialite-hand specimens, and sediment samples were simultaneously collected for mineralogical and geochemical analysis. The microbialite hand specimens were label as MB (n=6) and obtained using a scalpel. The sediment samples, which weighed ca. 100 g each, were denoted as S (n=4) and were collected using a scoop. Both sample types were carefully placed in plastic Ziplock bags for storage and transport. Additionally, we also collected discrete carbonate samples from various depositional settings in August 2022 for mineralogical identification. Specifically, we sampled carbonate mud (CM, n=2) and sand (CS, n=1) from Zone I, terrace deposits (CT, n=2) from Zones I and V, and carbonate crust on gravel (CC, n=2) from Zone VI.

For microbial community analysis, microbial mat samples were collected from the uppermost surface of the microbialites, and sediments in their proximity in August 2022. The sampled microbial mats that were denoted as ML(n=17) were immediately stored in a nucleic acid preservation solution (LifeGuard Soil Preservation solution, MoBio Laboratories, Carlsbad, CA, USA) and were placed in a freezer within 8h of sampling. They were then transported to the Istanbul Technical University –Geomicrobiology and Biogeochemistry Laboratory (ITU-GBL) where they were stored at -20°C until further analysis. The detailed descriptions of the samples subjected to microbial community analysis were provided in Table 1.

In August 2022, we conducted additional microbialite sampling at water depths of 5, 10 and 15 m in Zone I, Kocaadalar Burnu, also known as “White Island”, due to its extensive microbialite formations. The divers collected the samples in sterilized organic–free glass jars (ca. 200 ml) with wide mouths to collect fragments of the microbialites. The sample jars were filled with the hosting water to prevent exposure of the microbialites to atmospheric conditions.

After the collection, the sample jars (n=3) were stored at -21 °C until further examination and transported to the ITU-GBL. After thawing, the sample jars were placed into the safety cabinet and left under UV for 25 min to decontaminate the jars surfaces. The microbialite sample was then dissected with a pre-sterilized razor in the laminar flow based on color and textural differences for 16S rRNA amplicon sequencing. Four distinct layers were sampled and designated as MDL-1, MDL-2, MDL-3, and MDL-4 (Fig. 2). If possible, replicate samples (5 -10 gr), if possible, were stored for further geochemical and mineralogical analyses.

**Figure 2.**
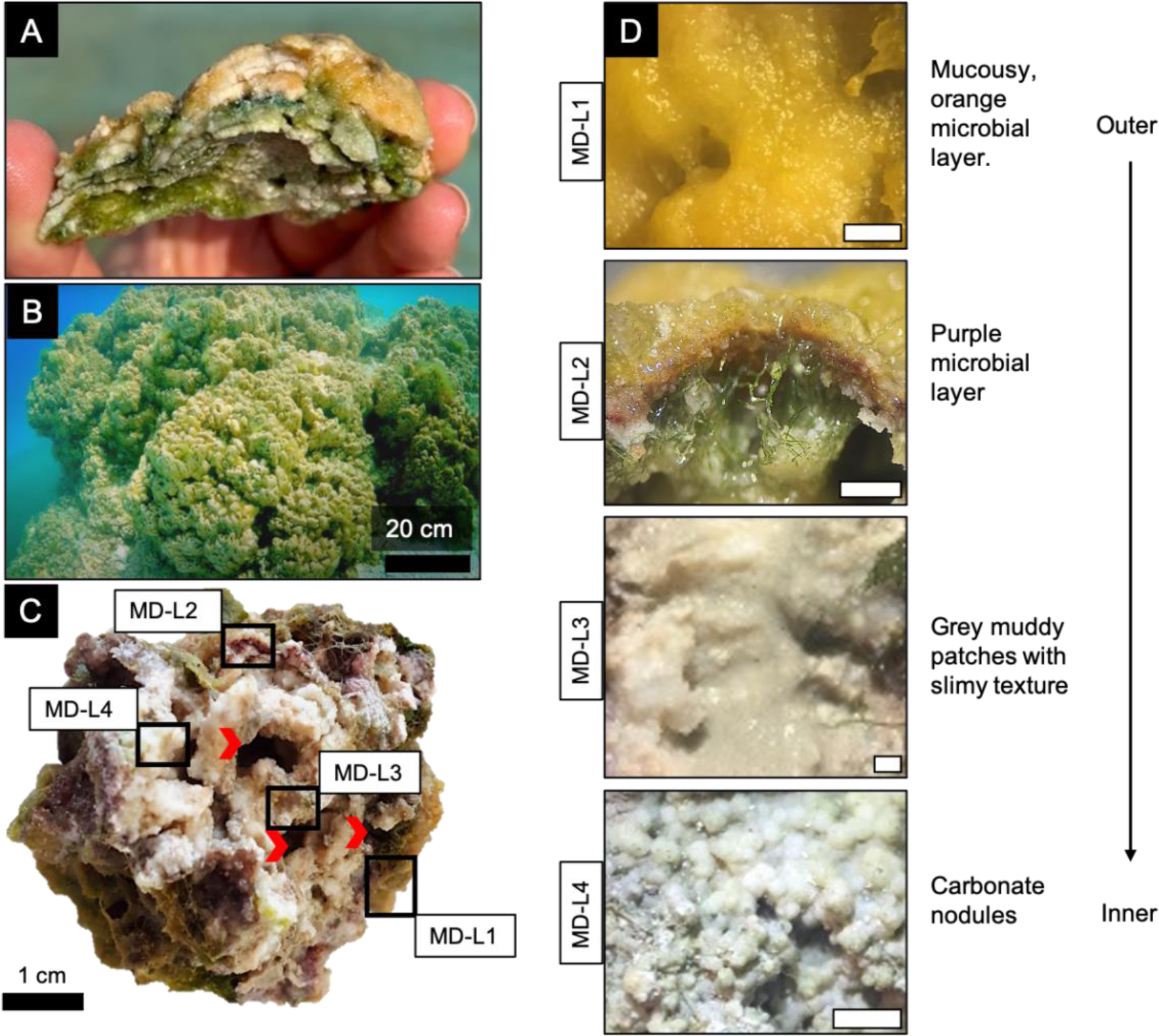
(A) Typical microbial mat development in the shallow microbialites. **(B)** General underwater image of the actively growing mini columnar thrombolitic microbialites at 15 m water depth in Zone I. **(C)** Close up view of the inner part of a hand specimen of thrombolitic microbialite in (B). Vuggy structures (red arrow) are common in the microbialites. **(D)** Representative close-up view of each dissected layer from outer to inner part of the sample in (C). White scale bar is 2mm.

### Geochemical Analysis

All water samples were analyzed at ACME Bureau Veritas Mineral Laboratories, Canada for elemental, cation and anions compositions (ACME, Analytical codes of AQ130, SO 200 and SO 001*). Alkalinity measurements were performed by colorimetric titration (MERCK Mcolortest^TM^) at the ITU-GBL laboratory. Simultaneously pH was measured using a glass pH electrode Unitrode. The alkalinity measurements had a detection limit and precision of 10 µeq L^-1^.

Major elemental chemistry of the microbialites and sediment samples subject to mineralogical analysis was determined by X-Ray Fluorescence (XRF) and Inductively Coupled Plasma Mass Spectrometer (ICP-MS; Perkin Elmer) at ACME laboratory (Analytical code of LF200), respectively.

The total organic carbon (TOC) content in the microbialites and surface sediments was analyzed with a Shimadzu TOC/TIC analyzer at ITU-Eastern Mediterranean Center for Oceanography and Limnology (EMCOL) Geochemistry Laboratory. A complete description of the methods used for these analyses can be found in Çagatay et al. (2015). The precision of the TOC analysis with this method is better than 2% at a 95% confidence level.

### Mineralogical and microscopic analysis

The mineralogical compositions of the microbialites (both sub/fossil and fresh), surface sediments and various carbonate samples were determined by X-Ray diffraction (XRD) at ITU. To obtain an average composition for each microbialite sample, including sub/fossil ones, a, ca. 10 mm^3^ sample was collected from the top, middle, and bottom of the sample. These randomly collected three samples were combined to create an overall average composite sample for each microbialite. The samples were hand crushed to a powdery size in an agate mortar and then thoroughly rinsed with de-ionized water to remove possible salts and low molecular weight organic particles. After rinsing, the samples were allowed to dry at room temperature. The dried samples were then collected on a silicon sample holder and analyzed using a Bruker diffractometer with CuKα radiation. Data collection was performed between 2 and 72 degrees with a total counting time of half an hour. The X’Pert HighScore Plus software was used for background subtraction, peak identification and matching with XRD patterns of reference compounds (US Institute of Standards and Technology, USA).

The microbial layer (ca. 2 mm) obtained from the actively growing microbialite at 15 m in Zone I was subjected to microscopic observations (Fig. 2D). To prepare the samples for microscopy, a fixation procedure was followed as described in Fowler (2011). This involved a three-day-long fixation process using sodium-glutaraldehyde and osmium tetra oxide, followed by gradual ethanol dry processes (30-50-70-90-99%). Scanning electron microscopy–energy dispersive spectroscopy (SEM-EDS) analytical techniques were used to provide a semi- quantitative elemental composition and spatial morphological analysis of the crystals in the layer, if present. Samples were coated with Au/Pd for 3 min to prevent charging and subsequently were analyzed by a Philips XL30 ESEM-FEG/EDAX system at Advanced Technologies Research Center, Bosphorus University.

### DNA extraction and 16S rRNA gene analysis

16S rRNA gene analysis was conducted to determine the prokaryotic diversity of the microbialites (n=12 + n=4 sub-sections) and sediments (n=5) in Lake Salda. Genomic DNA extraction from the samples was performed with the DNeasy PowerLyzer Power Soil Kit (QIAGEN, Hilden, Germany) following the manufacturer’s instructions. 16S rRNA coding regions of the extracted DNA were amplified by polymerase chain reaction (PCR) with universal EMP (Earth Microbiome Project) primer set including 515FB (GTGYCAGCMGCCGCGGTAA) and 806RB (GGACTACNVGGGTWTCTAAT) that encodes V4 region and targets nearly all bacterial and archaeal taxa (Caporaso *et al*., 2011).

PCR mixtures were prepared including 0.8X PCR Master Mix (PlatinumTM Hot Start PCR Master Mix (2X), Thermo Fisher Scientific, Invitrogen, Eugene OR, USA), 0.2 µM of each forward and reverse barcoded primers, 1.0 µL of template DNA, Phusion® High-Fidelity DNA polymerase (Thermo Fisher Scientific, Invitrogen, Eugene OR, USA), and DNA-free grade water up to a total reaction volume of 25 µL. PCR was performed starting with a single cycle of 94 ℃ for 3 min, 35 cycles of 94 ℃ for 45 s, 50 ℃ for 60 s, 72 ℃ for 90 s, and final single cycle of 72 ℃ for 10 min. Sequencing libraries were constructed with adaptor sequences and reaction conditions from TruSeq DNA LT Sample Prep Kit and sequenced on an Illumina iSeq 100 sequencer (Illumina, San Diego, CA, USA). The reads were filtered and trimmed to a minimum Phred quality score of 10. Following this step, the reads were merged, de-replicated, and clustered into operational taxonomic units (OTUs; 97% similarity) in QIIME2 (Bolyen *et al.,* 2019) using SILVA NR99 v.138 as the reference database (Quast *et al.,* 2013). Sequences derived from this study (16S rRNA gene) are available in GenBank under BioProject PRJNA1005563.

## RESULTS

### Water chemistry

The physicochemical and chemical parameters of the lake water were measured during the field campaigns in 2019, 2020, and 2021 (Table 2). Lake water contains high concentrations of Mg up to 350 mg /L (avg., 319 mg/L) but very low concentrations of Ca (avg., 0.59 mg/L) throughout both the wet (March) and dry (August) sampling seasons. The enrichment of Mg relative to Ca in the lake and surface waters suggests that it derived from the surface weathering of ultramafic rocks.

**Table 2.**
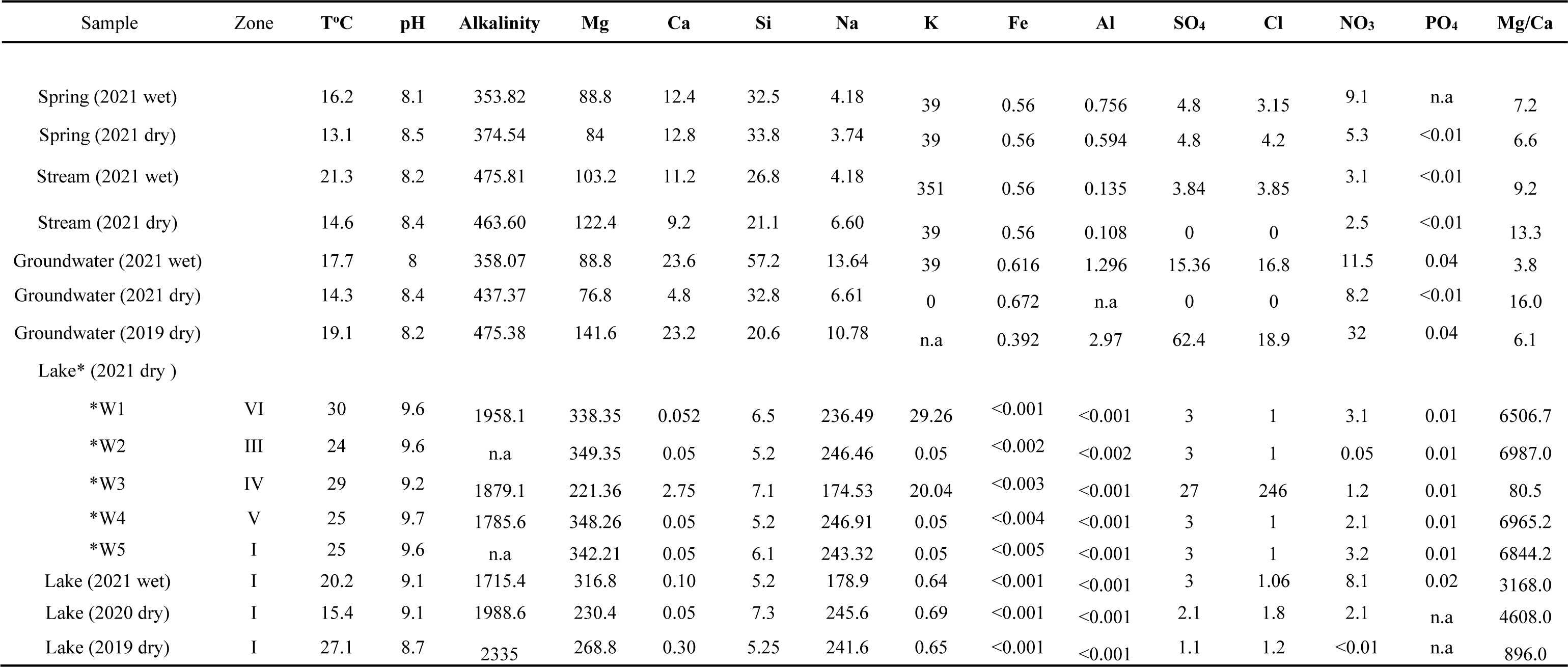
Geochemical and physicochemical characteristics of Lake Salda and near water sources (mg/L). Analyses not measured are listed as <u>“n.a”.</u>.

There are insignificant fluctuations in most water parameters between sampling campaigns, except for the alkalinity. Alkalinity ranges from 1785 to 2335 mg/L, with the highest values in the dry season. The average pH value of the lake water is of 9.6 ±0.2 in the dry season and 9.2±0.1 in the wet season. The concentrations of K (0.07 mg/L), Na (245.6 mg/L) and Cl (2 mg/L) show no significant changes among the sampling zones, except for Zone III, which has the highest K (20.04 mg/L), CI (246 mg/L), and sulfate (27 mg/L) concentrations. The lake water contains ca. 3 mg/L and 7 mg/L sulfate and silica, except for Zone III. The dissolved major constituents of lake water display the following sequence that can be considered as a general lake water characteristic without showing significant seasonal changes: Mg >> Na > K > Ca and HCO_3_+CO_3_ > Cl > SO_4_ > NO_3_ > PO_4_. The Mg/Ca molar ratios of lake water showed substantial differences among the zones. With an increase in chloride, sulfate, and potassium concentrations, Mg/Ca ratio decreased to the lowest ratio measured in the lake. Analysis of the lake water chemistry shows that the lake water is derived from meteoric water that mainly interacted with the ultramafic rocks prevailing both around the lake and lake basin. Unlike the lake water, the spring and stream waters that originate from serpentinized gravels and pebbles generally contain higher levels of silica, calcium, potassium, and sulfate. Groundwaters exhibit higher sulfate and chloride concentrations, along with ca. two-fold Mg and Ca concentrations compared to the spring and stream water (Table 2). Compared to fluvial water sources, the excess Mg in the lake water is attributed to the additional leaching of Mg by the meteoric water from ultramafic rocks, ultramafic-derived pebbles, and sand sized serpentinite-gabbro clasts in the alluvial fan deltas. These various size materials provide exceptionally large surface areas for enhanced meteoric water-rock interaction, releasing Mg into the environment.

### Geochemistry and Mineralogy of the microbialites

Mineralogical and geochemical data show that both fresh and fossil microbialites contain 35-40 % MgO, 0.74-1.43 % CaO, 0.07-1.0 % Fe_2_O_3_, 0.40-5.1% SiO_2._ No other elements exceed 1% in abundance (Table 3). They have a predominant content of 8–11 wt % TIC and 0.5–3 wt % TOC. The highest TOC content, exceeding 3 wt %, was measured in Zone I. There are no significant differences in the geochemical profile of the actively growing and fossil microbialites with two exceptions. The actively growing microbialites have slightly lower MgO (up to ca. 5 %) and higher SiO_2_ (up to 4 %) content, compared to fossil ones. Also, the SiO_2_ content of the microbialites is generally higher than that of the sediments. In terms of elemental sequence, the microbialites follow the typical pattern of Mg > Si=> Ca > Na > K > Fe > Al. The geochemical composition of the surface sediments, which mainly consist of hydromagnesite clasts (ranging from pebble to fine-sand) and gabbroic grains is quite consistent with that of the microbialites.

**Table 3.** Geochemical composition (wt.%) of the fresh microbialites, sub-fossil microbialites (MB) and sediments (S).

| Sample | Zone | Condition | MgO | SiO<sub>2</sub> | CaO | Na<sub>2</sub>O | K<sub>2</sub>O | Fe<sub>2</sub>O<sub>3</sub> | Al<sub>2</sub>O<sub>3</sub> | LOI | TC | TIC | TOC |
| --- | --- | --- | --- | --- | --- | --- | --- | --- | --- | --- | --- | --- | --- |
| MB1<br>(Fig. 8E) | I | Fresh<br>(Shallow < 1m) | 38.59 | 0.40 | 0.97 | 0.06 | 0.02 | 0.06 | 0.05 | 59.2 | 11.25 | 9.84 | 1.41 |
| MB2<br>(Fig. 2C) | I | Fresh (Deep >15m) | 38.10 | 0.72 | 1.17 | 0.04 | 0.02 | 0.11 | 0.10 | 59.1 | 11.47 | 9.93 | 1.54 |
| MB3 | IV | Sub-fossil | 38.12 | 3.06 | 1.17 | 0.37 | 0.05 | 0.52 | 0.23 | 55.7 | n.d. | n.d. | n.d. |
| MB4 | VI | Sub-fossil | 37.77 | 3.94 | 1.43 | 0.24 | 0.04 | 0.78 | 0.31 | 54.7 | n.d. | n.d. | n.d. |
| MB5 | VI | Sub-fossil | 36.98 | 3.05 | 0.74 | 0.35 | 0.09 | 0.56 | 0.29 | 57.2 | 10.83 | 9.25 | 1.58 |
| MB6<br>(Fig. 8B,C) | VI | Sub-fossil | 36.75 | 4.60 | 0.90 | 0.10 | 0.04 | 1.00 | 0.45 | 55.3 | 10.63 | 8.48 | 2.15 |
| S1 | I | White beach deposit (pebble) | 40.60 | 0.31 | 0.86 | 0.01 | 0.00 | 0.07 | 0.04 | 57.4 | 10.98 | 10.57 | 0.41 |
| S2 | V | Terrace deposit | 39.25 | 1.66 | 1.40 | 0.02 | 0.02 | 0.24 | 0.18 | 56.5 | 11.12 | 10.25 | 0.87 |
| S3 | I | White beach deposit (sand) | 39.72 | 0.55 | 1.01 | 0.02 | 0.00 | 0.07 | 0.04 | 57.9 | n.d. | n.d. | n.d. |
| S4 | I | Shallow lake bottom sand | 35.87 | 5.05 | 0.83 | 0.07 | 0.07 | 0.83 | 0.67 | 55.9 | 11.06 | 7.66 | 3.40 |

Hydromagnesite was the dominant carbonate mineral in all sampled microbialites. Similarly, mineralogical compositions of various carbonates such as mud (CM) and sand (CS) deposited on the shoreline and carbonate crust (CC) on gravel and pebbles around the lake also comprised mainly of hydromagnesite (Fig.3). In addition to this, quartz (SiO_2_) is present in both carbonate crust and mud while nesquehonite (Mg(CO_3_)·3H_2_O) is only found in the mud. Aragonite is present in both carbonate terraces (CT) and in the mid-section of 15 meters submerged microbialite sample (MDL4) (Fig. 3). The mineralogical composition of both fresh (MDL4) and fossil (CT) microbialites aligns well with their elemental abundances. These microbialites mainly consist of hydromagnesite with a minor amount of aragonite as indicated by the relative peak intensity. One of the carbonate terrace samples also contains diopside (MgCaSi_2_O_6_) and hercynite (Al_2_FeO_4_) minerals.

**Figure 3.**
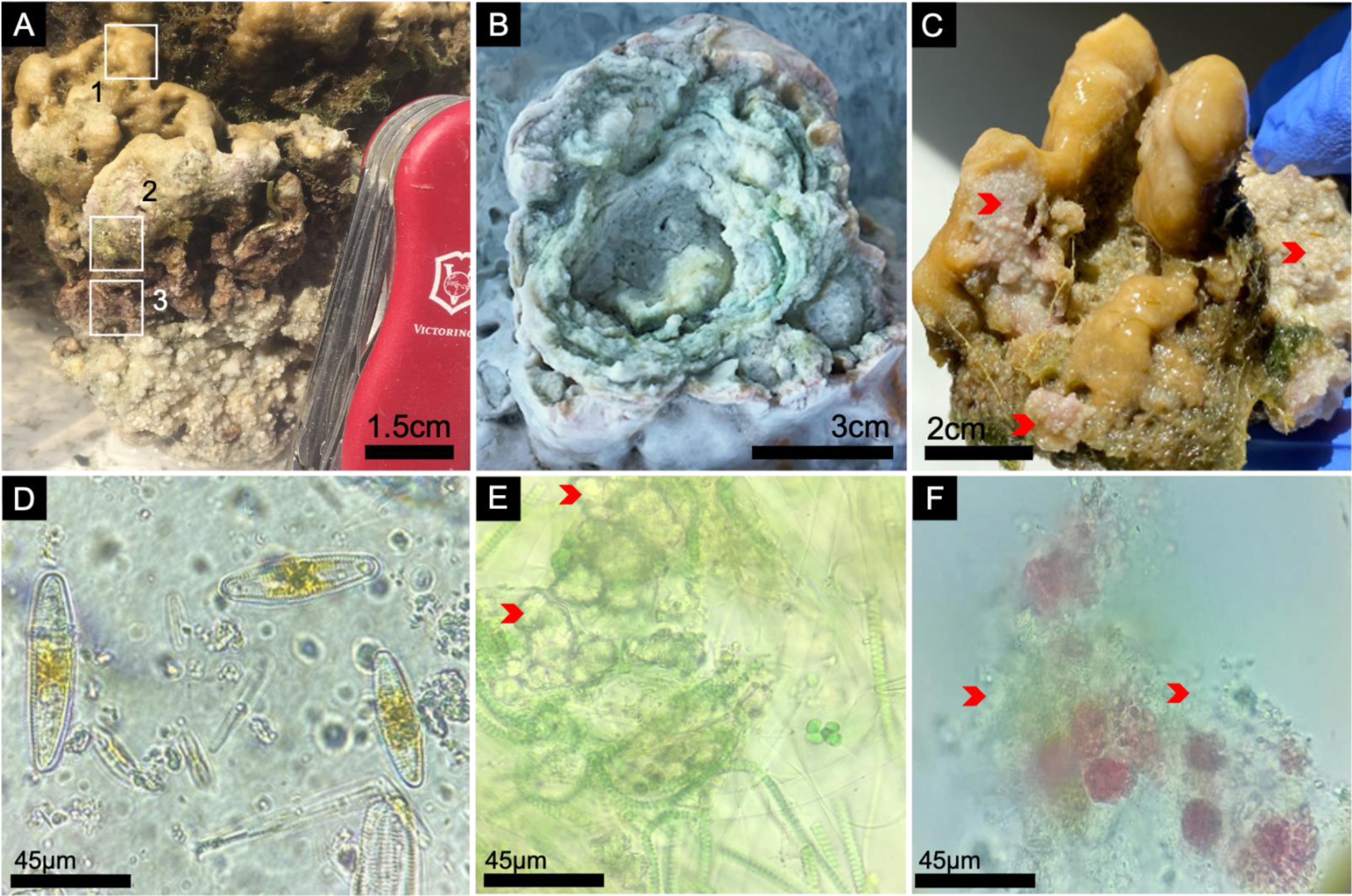
X-Ray Diffraction patterns of various carbonate structures, microbialites and microbial layers from Lake Salda. Carbonate mud (CM, n=2) from Zone I; carbonate crust on gravel (CC, n=2) from Zone VI; terrace deposits (CT, n=2) from Zone V; upper most surface (MDL1), and mid section (MDL4) of the mini columnar thrombolitic microbialite from 15 m water depth (see Fig. 2) and carbonate sand (CS, n=1) from Zone I. H: Hydromagnesite, Q: Quartz, Hc: Hercynite, D: Diopside, A: Aragonite, P: Palygorskite, N: Nesquehonite.

Additional XRD analysis was conducted with individual samples being collected from the columnar linked microbialites that grow at 15 m water depth in Zone I (Fig.2 B, C). The uppermost surface of the sample was characterized by yellow/orange-colored, organic rich mucilaginous layer rich in diatoms (MLD1, Fig. 2C&D). It also contained Mg and Si as palygorskite (Mg,Al)_2_Si_4_O_10_(OH) along with hydromagnesite mineral (Fig. 3). Although various amount of silica was detected within the microbialites, palygorskite mineral was only detected in the top mm diatom- rich surface section of this deeply growing microbialite (Fig. 2D). Furthermore, unlike general mineralogical composition of the microbialites, no aragonite was present in this sample. In contrast, as suggested by the peak intensity, relatively higher aragonite with hydromagnesite was present in the microbial layer that contains the carbonate nodules (Fig. 2D). Although aragonite was consistently detected within the microbialites (both fresh and sub/fossils) in the lake, it is first time that aragonite associated with a particular microbial layer within the microbial mat is identified (Figs. 2 and 3).

### Microscopic characterization of microbial mat

Microbial mats of the microbialites in Lake Salda do not always show successive layering and the thickness of each layer, whenever present, varied from site to site in the lake (Fig. 2A). Here, we described typical characteristics of microbial mat of the microbialites. The microbial mats generally have a mucilaginous dark –orange to yellow-greenish colored surface layer, which is typically a few millimeters thick but can occasionally reach centimeter-scale thickness. This surface layer is consistently present in both shallow and deeply growing microbialites (Fig. 2A,D and Fig. 4A,C). Beneath the surface layer, there are lithified endolithic layers that can vary in color, ranging from lighter to darker green. These green-colored endolithic layers are often well observed within the finger-shaped mini columns in the shallow growing microbialites, which are found at around 1-1.5 meters of water depth (Fig. 2A). These endolithic layers can also be observed in the deeply growing microbialites (Fig. 4C). Below the green-colored layers, there are typically purple- and pink-colored layers, particularly within the deeply growing microbialites. These layers contribute to the overall composition and appearance of the microbialites.

**Figure 4.**
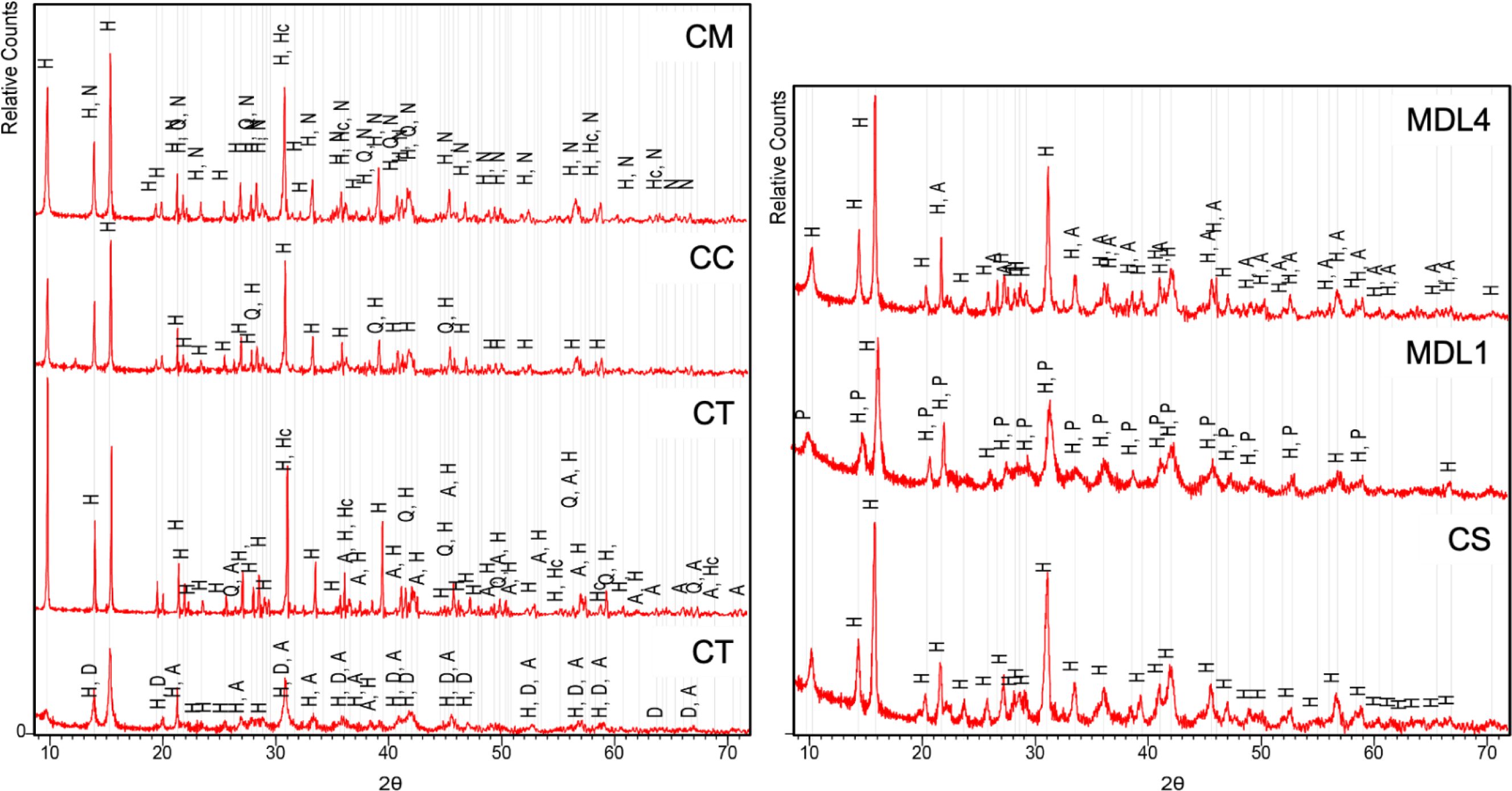
Field views of microbialites **(A-C)** and light microscopy images **(D-F)** of microbial mats collected from Lake Salda. **(A)** A typical microbial mat of the deeply growing thrombolitic microbialites (> 10 m) in Zone I. **(B)** Microbial mat and internal structure of stromatolitic thrombolites in Zone VI. **(C)** A close view of the finger shape mini-columns from the deeply growing thrombolites in Zone I (> 10 m), note the internal grainy structure. **(D, E, F)** Light microscopy images of layer 1; 2 and 3 in A, respectively; (red arrow indicates carbonate precipitation).

Light microscopy images of the top mm orange-colored layer showed an extensive presence of diatoms with filamentous and coccus-shaped cyanobacteria (Fig. 4D). The green endolithic microbial layer is dominated by filamentous cyanobacteria and carbonate precipitation (Fig. 4E) while the pinkish and purple layer revealed aggregated microbial community within the precipitation (Fig. 4F).

Consistent with the light microscopy images SEM image of the orange-colored microbial layer, in particular, at a meter’s depth (> 10 m), showed a remarkable of diversity diatoms in different morphologies. These diatom species *(Cymbella* sp., *Gomphonema* sp., *Pinnularia* sp. *Gomphonema* sp., *Synedra* sp. *Eunotioid* sp., *Epithemia adnate, and Navicula recens)* were previously identified by Balci *et al*. (2018). The diatom, filamentous cyanobacteria and mineral aggregates form a porous network (Fig. 5A). Mineral aggregates within the organic matrix –exopolymeric substances (EPSs) are generally sub/spherical to slightly elongated and are usually 10-50 µm in diameter (Fig. 5B). Carbonates are also found as bigger clusters embedded in the EPS matrix (Fig. 5B). Some diatoms can be observed inside the mineral aggregates as well as at the outer edge. They are usually well preserved, and significant fragmentation or corrosion features of frustules are rare (Fig. 5A,B).

**Figure. 5.**
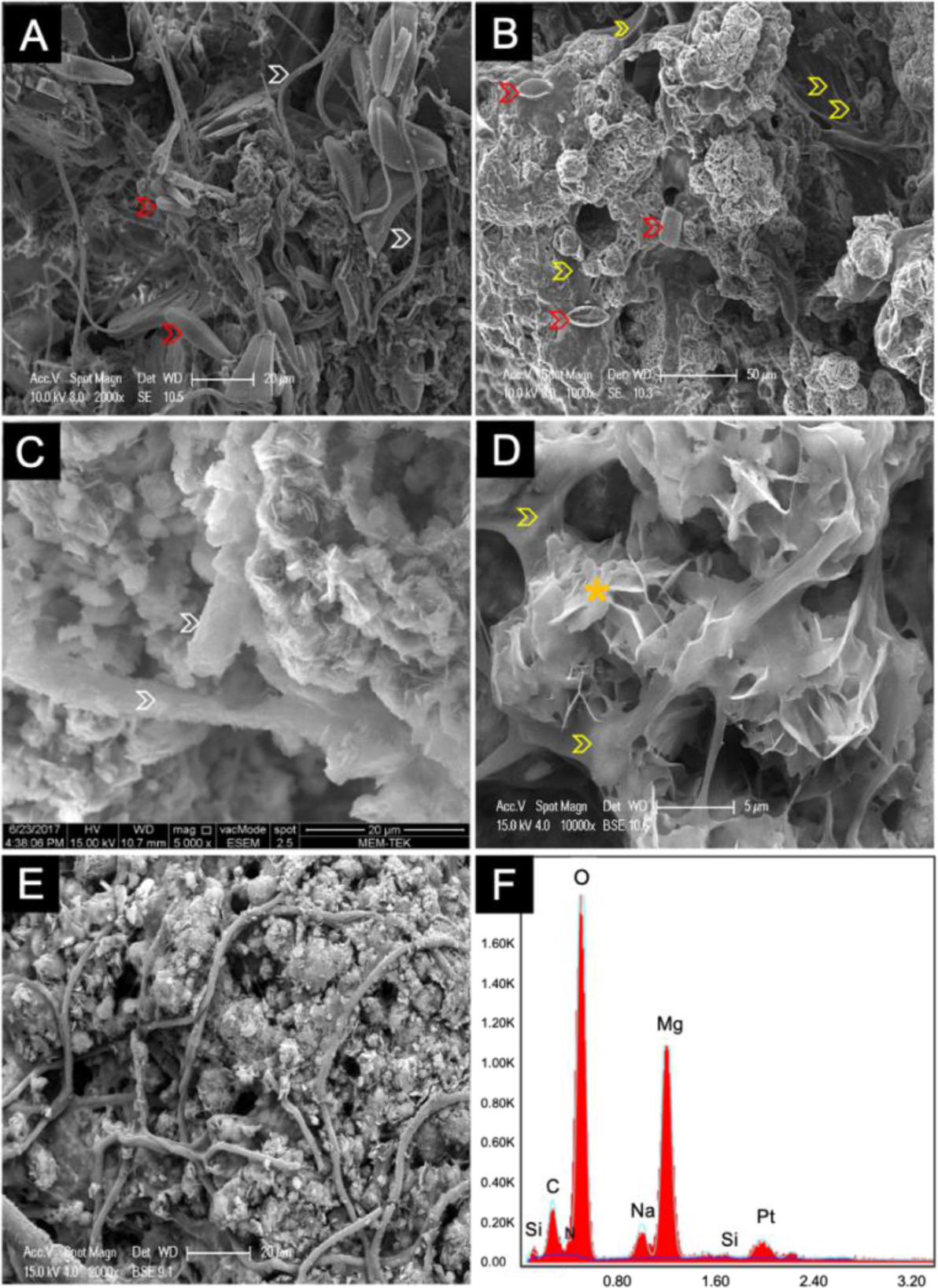
Scanning electron microscope images of the microbial layers of microbialites. **(A)** Diatom-filament –mineral aggregates identified in the orange-colored layer (see Fig. 4A,1). **(B)** Sub/spherical and blocky carbonates within the EPS rich zones. **(C)** Carbonates and carbonated filaments within the green colored layer (see Fig. 4B). **(D)** Plate and blade like crystals, characteristic of hydromagnesite, found in the green section of stromatolitic thrombolites. **(E)** Non-carbonated filaments in the stromatolitic thrombolite. Remains of EPS material are indicated by yellow arrows while diatoms and filaments are indicated by red and white arrows, respectively. **(F)** EDS spectra of the region marked by the yellow star in C.

SEM images of the endolithic greenish colored layer showed a copious amount of precipitation that is generally associated with the filaments and organic matrix-EPS (Fig. 5C, D). Similarly, SEM images, in particular those from stromatolitic-thrombolites in Zone VI (Fig. 4B), showed that some of the filaments and EPS are covered with the aggregated carbonates (Fig. 5C,D). The carbonate material that covers the filaments in the mineralized parts of the matrix is represented by mainly plate-like, radially divergent crystals that typically formed sub/spherical clusters (Fig. 5C,D) but some filaments appear to be not carbonated and most likely provide structural stability (Fig. 5E). Some of the EPS material also exhibited bubbling with HCl, indicating that these polymeric substances might also have carbonate material present in them. In contrast to the greenish colored layer with Mg, C and O content, EDS spectra of the yellowish-orange microbial layer, revealed the presence of C, O, Mg, in addition to Ca, and Si (Fig. 5F).

### Morphological diversity and distribution of microbialites

Lake Salda’s hydromagnesite build-ups were initially named as stromatolites (Braithwaite and Zedef, 1994; 1996; Shirokova et al., 2011). Here, we embrace the umbrella term “microbialites” since various microbialite structures as described below are present in the lake. The morphological diversity of the microbialites is, for the first time, described in detail and several previously unexplored morphotypes of the microbialites were identified. A representative example for each type along the entire perimeter of the lake is presented at the macro (Fig. 6) and meso scale (Figs.7 and 8). Accordingly, six major zones that host various contemporary hydromagnesite build-ups were described along the lake.

**Figure 6.**
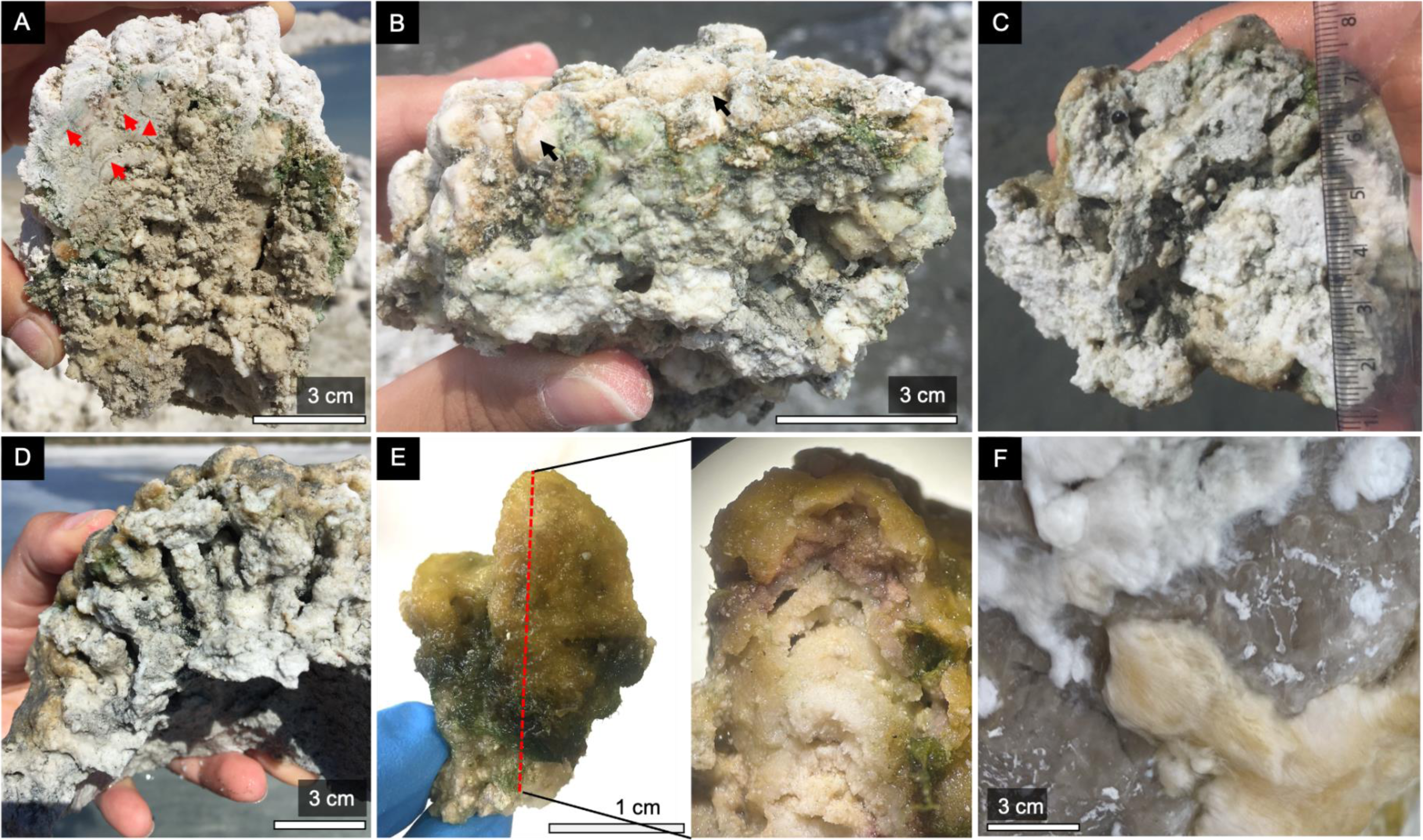
Identified morphotypes of the microbialites in Lake Salda. **(A)** Shallow-growing dome-shaped microbialites with cauliflower morphology of various sizes and fossil microbialites (white islands) in Zone I. **(B)** A dome-shaped radially growing microbialite with colloform surface texture in Zone III-IV (<1 m water depth). **(C)** The deeply growing microbialites with mini-column structures on the top surface in Zone I (> 10 m water depth) (see Fig. 7C). **(D)** Recently exposed sub/fossil microbialites with the mini column structures on the top in Zone II and **(E)** ridge morphology with cerebroid surface texture in Zone VI. **(F)** Stacked-domed-shaped microbialites with smooth surface texture and bulbous growth structure in Zone VI. **(G)** Laterally growing microbialites with hemispheroid macrostructures (Image from Brad Garczynski). **(H, I)** Microbial pavements in Zone I and III.

The contemporary hydromagnesite build-ups with thrombolitic structures commonly occur along the shore and at varying water depth of the lake (< 20 m) (Fig.6 A,B,C). Accordingly, the half-domed or dome-shaped thrombolitic build-ups with cauliflower morphology and colloform surface texture are particularly abundant in Zone I, II and IV (Fig.6 A,B,C). They are a few cm to meter scale in diameter and accrete sub-aqueously at a water depth of 1 or < 1 meter (Fig. 6A and B). In Zone I, exposed build-ups even form an island (White Island), a structure ca. 10 and 20 m in diameter and 4-5 m in height (Fig. 6A). Thrombolites with columnar-linked morphology at a water depth of ca. 15 m are recently identified (Fig. 6C). Compared to the radially growing shallow build-ups with domed-shaped morphology the upper surface of these deeply and vertically growing thrombolitic structures (< 15 m of water depth) are covered by the finger-shape mini columns (a few cm in size) which are typically interconnected (Fig. 6C). Recently exposed subfossil thrombolitic structures in Zone II have well-developed and preserved, finger shape mini-columns on the uppermost surface (Fig. 6D). Hydromagnesite build-ups with cerebroid surface texture exist as pavement like structure particularly in Zones III, IV and VI (Fig. 6E). Stacked-domed shaped thrombolites with typical bulbous morphology are most clearly observed in Zone VI (Fig. 6F). Moreover, in Zones IV and V (ca. ∼2 m water depth), hydromagnesite build-ups (∼25-50 cm in relief) typically exist as laterally growing hemispheroid macrostructures and the finger-shaped mini columns, as observed in Zone I, cover the upper surface of these hemispheroid structures (Fig. 6G). Microbial pavements with smooth surface texture are excessively present in Zones I and III (Fig. 6H,I). The upper surface of the pavements is completely covered by a few mm or cm scale smooth, whitish grey colored fenestrated microbial layer (Fig. 6I).

As observed in the macro morphology, at mesoscale, a remarkable diversity also exists within the identified microbialites (Fig. 7 and 8). It is here worth noting that all the microbialites with cauliflower and calcareous knobs top described here, possess the same overall external morphological appearance, thus dome or half-domed and linked columnar cauliflower morphologies. Near-shore cauliflower top domed or half-domed shaped microbialites exhibited continuous fine laminations (ca. 4 cm), at sub/millimeter scale, and growing upwards, on the uppermost surface compared to the granular bottom portion (Fig. 7A). In the same zone, calcareous knobs top microbialites have occasional lamination but instead have a few bulging carbonate knobs on the uppermost surface, resembling smaller mounds (Fig. 7B). These stromatolitic/thrombolitic structures are underlined by continues and massif carbonate layers (Fig. 7B). Additionally, the microbialites in Zone IV also showed stromatolitic thrombolite structure with granular internal structure with a knobby surface (Fig. 7C). Both cauliflower-top and calcareous-top microbialites with different internal growth structures were found almost at similar depths of 70–100 cm. Despite the similar external morphological appearance to those in Zone I and II, another internal growth structure was identified at a depth of ca. 50 cm in Zones III and IV, called as the branching or dentrolitic growth structure (Fig. 7D). The uppermost surface of these newly identified microbialites resembles to those in Zone I, representing several bulging carbonate knobs on the uppermost surface. In contrast to the shallow- growing microbialites, the finger-shaped mini columnar microbialites identified at a depth of ca. 15 m in Zone I exhibits a granular texture with visible sub-mm thick lamination (Fig. 7E). An orange and mucilaginous layer with sporadic algal material is present between the columns and on the uppermost surface (Fig. 7E). Occasionally, carbonate precipitates embedded within the white to orange-colored tufty biofilm covering rock surfaces on the shoreline can be observed, most commonly in Zone V (Fig. 7F).

**Figure 7.**
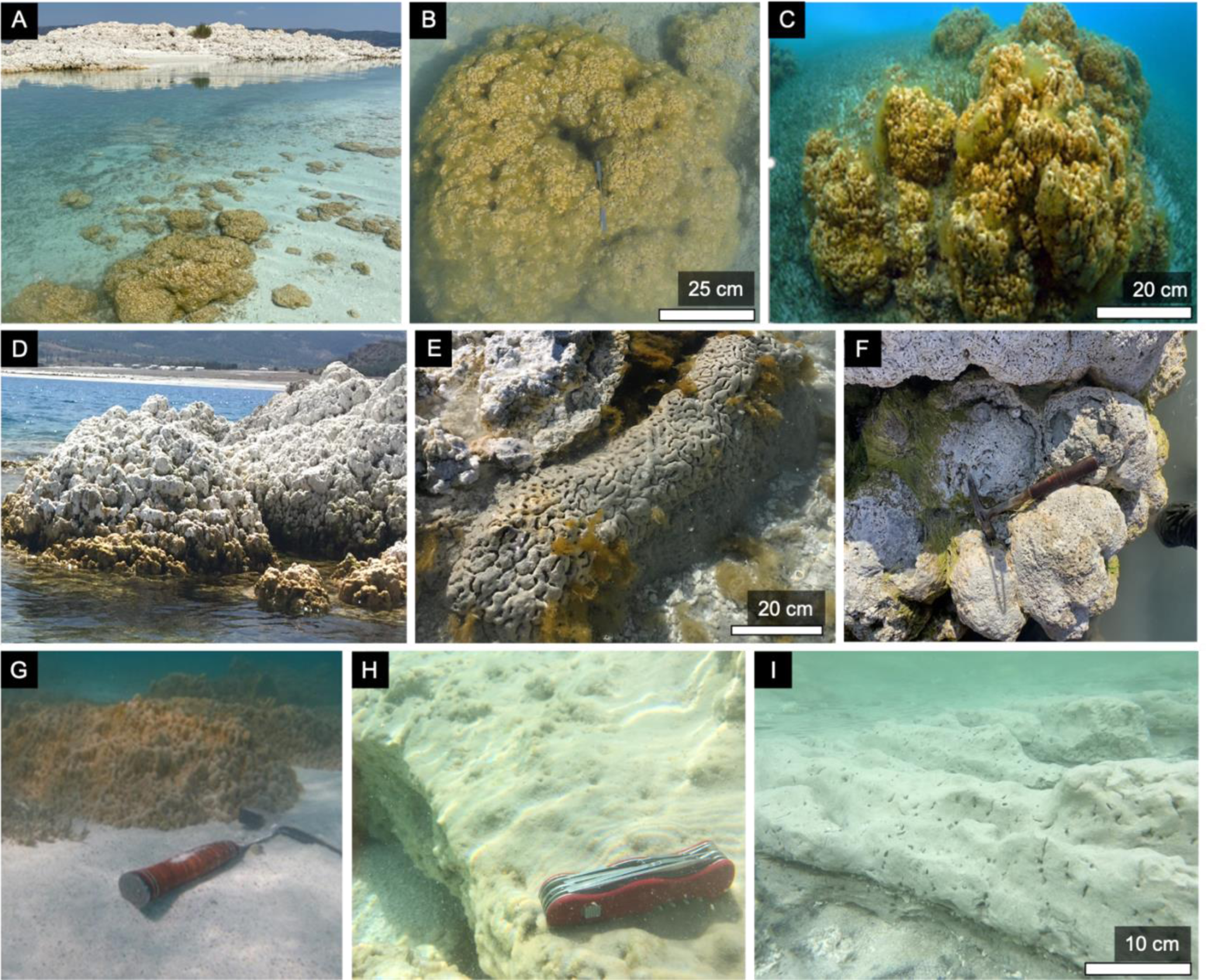
Various mesoscale internal structures identified in the lake. **(A)** Stromatolitic layers (red arrow) with cauliflower top in Zone I. **(B)** Thrombolitic growth structure with calcareous knob top (black arrow). **(C)** Thrombolitic microbialite with massif carbonate (see Fig. 6C). **(D)** Thrombolitic microbialite with branching growth structure. **(E)** A mini column with internal layers that grows on the top of the deep microbialites in Zone I (> 10 m water depth). **(F)** White and yellow biofilms with tufty texture covering a rock surface (Zone V).

**Figure 8.**
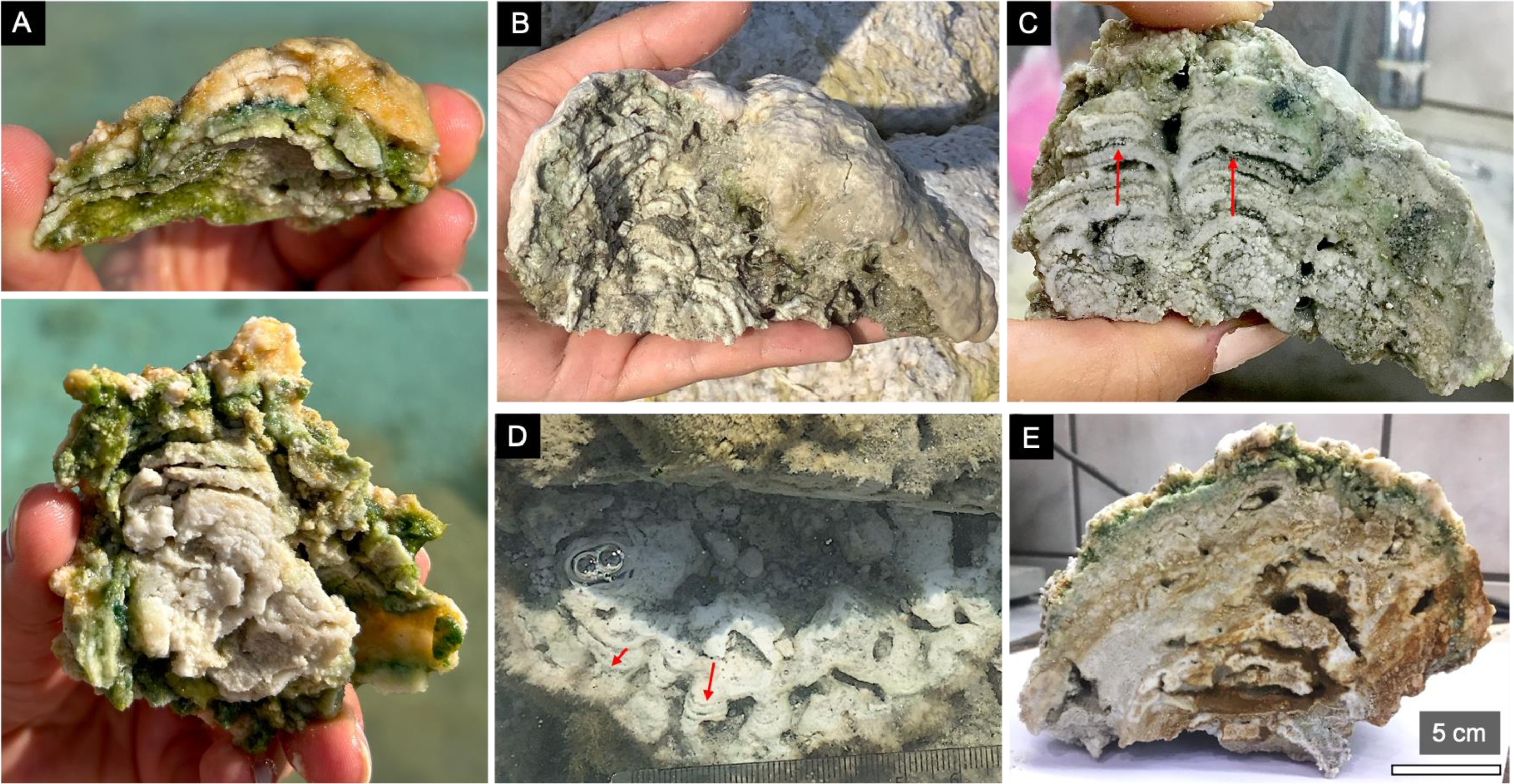
(A) Vertical cross section of a typical microbial mat with smooth surface texture and internal stromatolitic structure (above) and its inner core (below) in Zone I. **(B)** Sub-fossil counterpart of stromatolitic structure. **(C)** Internal structure of (B) with clotted thrombolitic section at the bottom and stromatolitic layers at the top. **(D)** Field image of the radially grown stromatolitic thrombolites in Zone VI, note the outward growing columnar structures (red arrow). **(E)** Vertical cross section of a hand specimen of diagenetic microbialite (Zone I).

At the meso scale, a well-developed stromatolitic microbialite sample from Zone I demonstrates typical layering structures of a shallow (ca. <1 m water depth) grown microbialite for the lake (Fig. 8A). In this stromatolitic structure, as always, orange mucilaginous top layer overlies on a dark pine green and bright pistachio green layers (Fig. 8A-top), reaching a massive carbonate core (Fig. 8A-bottom). Similarly, at the meso scale, stromatolitic thrombolites are commonly observed within the stacked domed shapes with smooth surface texture in Zone VI (Fig. 8B (see Fig. 6F for macro scale)). The half-sectioned stacked-domed shape revealed a thrombolitic base and stromatolitic layers at sub/millimeter scale growing radially and outwards (Fig. 8B,C). The radially grown lamination can be found in a few mm sizes, forming a cm-scale microbialite. Both stromatolitic and thrombolitic sections can be even seen in one structure (Fig. 8C). Compared to the dome shaped heads, elongated build-ups that show more ridge-like morphology and cerebroid surface texture identified in Zone III and VI show different internal meso structure (Fig. 6E, Fig. 8D). Mini columns with heights ranging from 2 to 5 cm are generally branching at the center and grow towards to the outward, creating radially growing stromatolitic structures (Fig. 8D).

It is noteworthy that while some thrombolites have well-preserved mesoclots that likely to reflect microbial growth in the lake, other thrombolites lack distinct mesoscale structures. This complicates identifying thrombolite growth structures and distinguishing them from other post-depositional features that may also have a patchy distribution in the lake. For example, a dome-shaped build-up has stratified features with a few mm- to cm-scale irregular cavities along with a mesoclotted or patchy fabric that may be defined as thrombolitic structure (Fig. 8E).

### Microbial diversity

A comprehensive cultivation-independent microbial diversity analysis was conducted on the microbial mats (ML, n=17) and sediment (MS=5) samples collected from different zones (I, III, IV, V and VI) and water depths (MD-L; 5 m, 10 m and 15 m). The description of the samples subjected to 16S rRNA analysis was provided in Table 1 while the sequencing data are summarized in Table 4.

**Table 4.** Summary of the bacterial 16S rRNA gene diversity analysis by the Illumina Hiseq Sequencing method.

|  | Sample | Sequences | OTUs | Shannon | Chao1 |
| --- | --- | --- | --- | --- | --- |
| Zone I | ML1-I | 8210 | 183 | 3.140 ± 0.02 | 59 |
|  | ML2-I | 135789 | 495 | 5.086 ± 0.02 | 361 |
|  | ML3-I | 15687 | 229 | 3.068 ± 0.01 | 70 |
|  | ML4-I | 9171 | 230 | 3.327 ± 0.02 | 76 |
|  | ML5-I | 9644 | 234 | 2.955 ± 0.01 | 72 |
|  | ML6-I | 3542 | 138 | 3.069 ± 0.01 | 50 |
|  | ML7-I | 109786 | 313 | 3.991 ± 0.02 | 190 |
|  | ML8-I | 60979 | 444 | 4.675 ± 0.02 | 356 |
|  | MLD1 | 16834 | 98 | 2.739 ± 0.01 | 41 |
|  | MLD2 | 7684 | 161 | 3.415 ± 0.02 | 101 |
|  | MDL3 | 12978 | 172 | 2.203 ± 0.01 | 63 |
|  | MDL4 | 13264 | 305 | 2.711 ± 0.01 | 60 |
|  | MS1-I | 19837 | 189 | 3.670 ± 0.02 | 133 |
|  | MS2-I | 7462 | 192 | 3.250 ± 0.02 | 65 |
|  | MS3-I | 8603 | 183 | 3.290 ± 0.02 | 78 |
|  | MS4-I | 9398 | 267 | 3.690 ± 0.01 | 117 |
| Zone III | ML1-III | 7778 | 179 | 3.713 ± 0.01 | 132 |
| Zone IV | ML1-IV | 17839 | 307 | 4.065 ± 0.02 | 166 |
| Zone IV | ML2-IV | 24594 | 272 | 3.993 ± 0.02 | 145 |
| Zone V | ML1-V | 119769 | 435 | 4.865 ± 0.02 | 321 |
| Zone VI | MS1-VI | 3541 | 165 | 3.755 ± 0.01 | 90 |

Bacterial sequences accounted for 99.6% of the total sequences, while archaeal sequences represented only a minor portion (0.4%) of the microbial community. Thirteen major bacterial phyla and one archaeal phylum were detected (Fig. 9). The dominant bacterial phyla were Proteobacteria, Chloroflexi, Firmicutes, and Bacteroidota, which together constituted approximately 80% of the microbial population in the microbial layers across different locations. Other phyla such as Verrucomicrobiota, Cyanobacteria, Planctomycetota, and Actinobacteriota were also present but at lower abundances. Bathyarchaeota was the only archaeal phylum identified in the entire lake.

**Figure 9.**
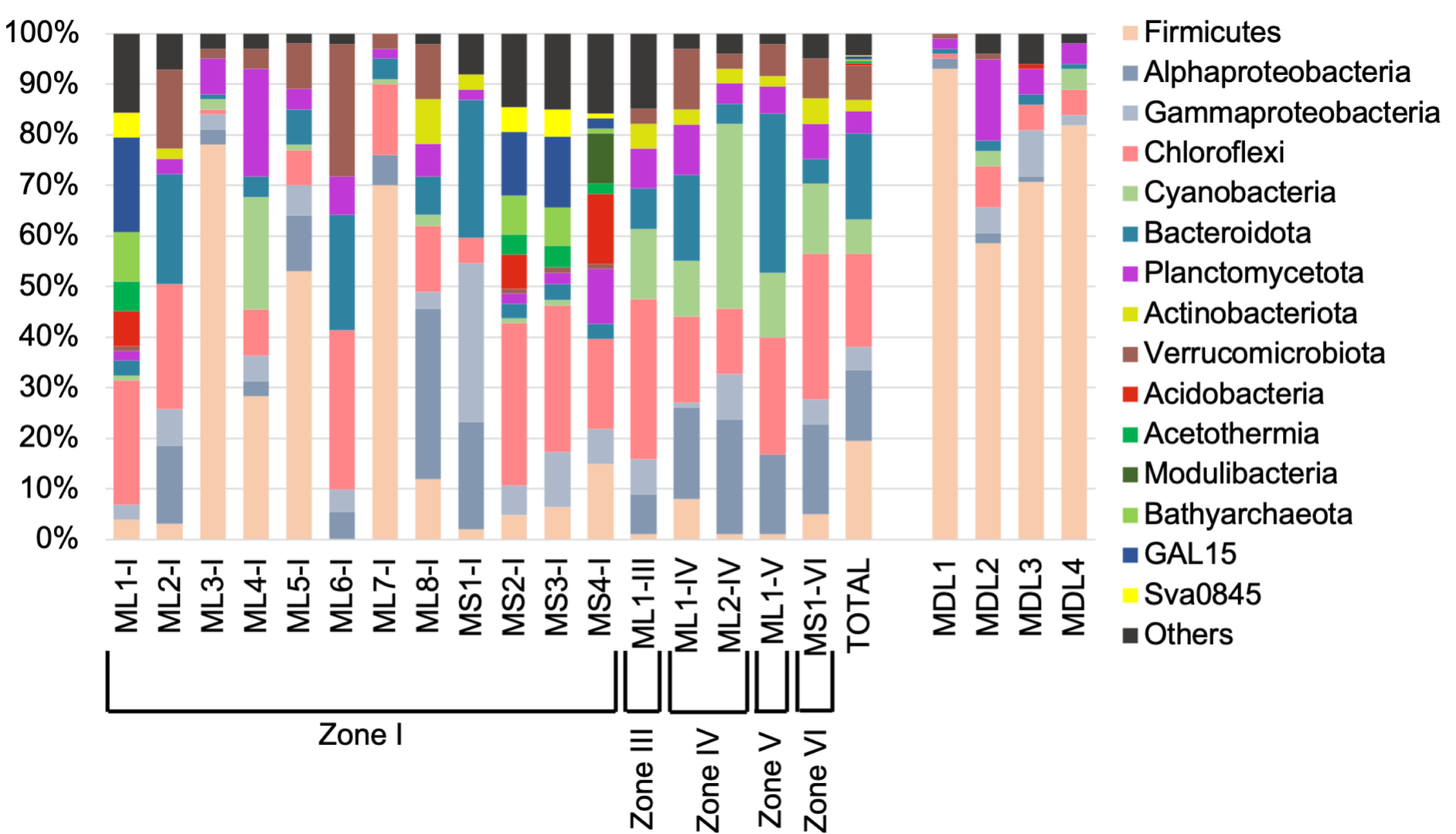
Relative percentage abundance of major bacterial taxa, as detected by 16S rRNA analysis, in the shallow (ML) and deeply growing (MDL) microbialites and the lake shore sediments (MS).

In Zone I and II, deeper thrombolitic microbialites had a higher percentage of Firmicutes phylum compared to the shallow ones. Chloroflexi constituted approximately 10% of the bacterial population in both the deeper and shallow water thrombolites, while Cyanobacteria constituted only a small proportion except for one sample (ML4-I). In Zones III, IV, V, and VI, Chloroflexi and Cyanobacteria accounted for approximately half of the microbial populations in the stacked domed shape microbialites, mainly stromatolitic thrombolites, as well as microbialites with dendritic growth structures. Gammaproteobacteria, Bacteroidota, and Verrucomicrobiota were also present at significant abundances in these zones. Sediments had higher percentages of Chloroflexi, Bathyarcheaota, and GAL-15 clade.

As indicated by the phylogenetic data, the dominant genera detected in the microbial mat samples show significant spatial differences. For example, genera of A4b and SBR1031 within the phylum Chloroflexi, and *Coleofasciculus, Leptolyngbya, Nodosilinea,* and *Phormidium* genera within the phylum of Cyanobacteria were detected in high abundance in the stromatolitic thrombolites in all zones (ML1-III, ML1-IV, ML2-IV, MS1-VI, ML1-V, ML6-I, ML2-I) except Zone I (Fig. 10). Additionally, *Luteolibacter* (Verrucomicrobiota) was the other genus identified in high abundance.

**Figure 10.**
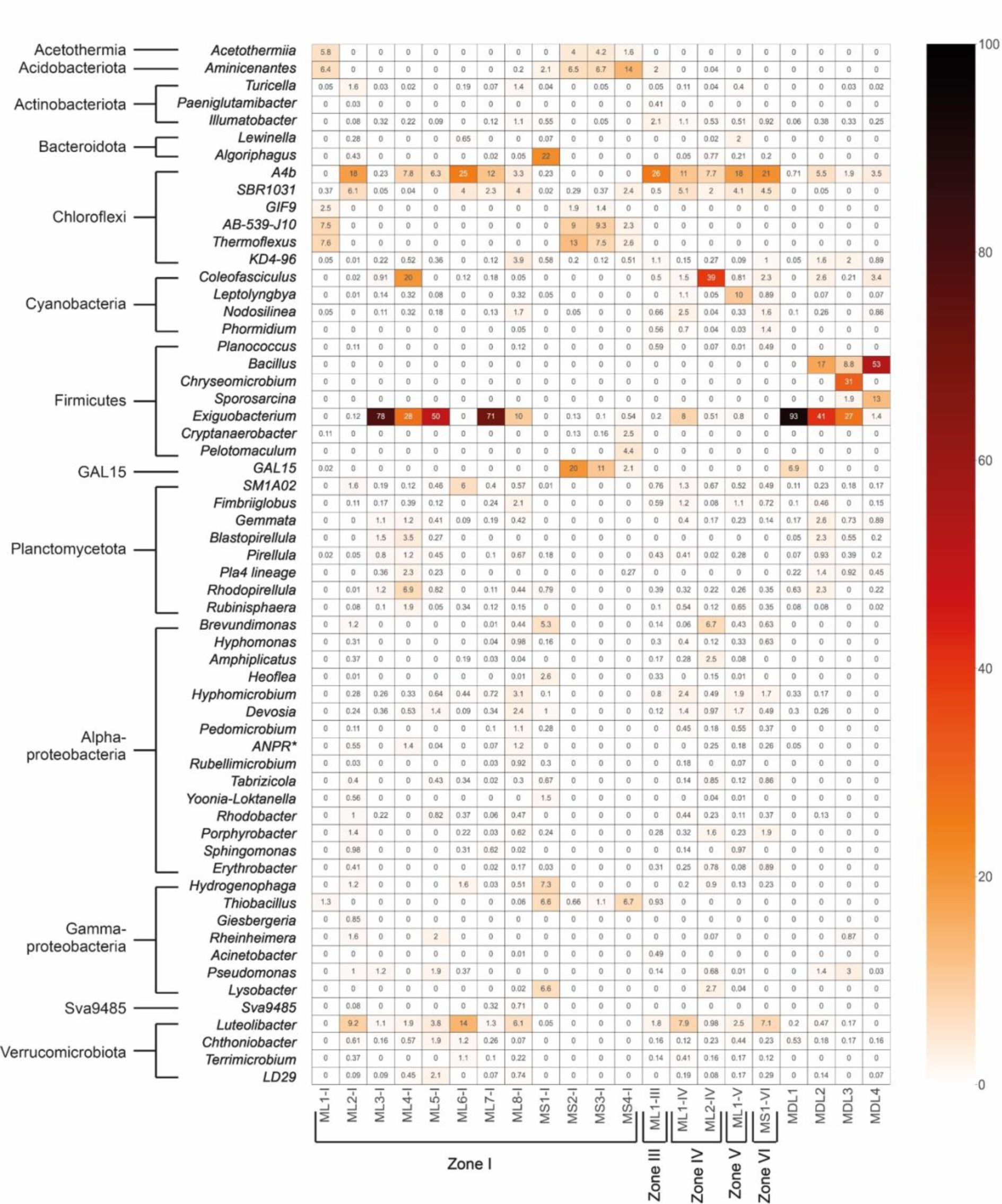
A heatmap showing the relative abundance (%) of the most dominant bacterial genera detected in the shallow (ML) and deeply growing (MDL) microbialites and the lake shore sediments (MS).

Thrombolites in Zone I, in particular, those of the deeper thrombolites (ML7-I, ML3-I, ML4-I, ML5-I) are dominated by Firmicutes genus *Exiguobacterium*, comprising 28-93% of the microbial community. The highest abundance of *Exiguobacterium* was observed in the uppermost orange-colored microbial layer of the deeper thrombolites at a depth of 10 m (MDL- 1), reaching 93%, as well as in the shallow cauliflower top and calcareous knobs top microbialites in Zone I. In Zone I, only genera of *Coleofasciculus* within the phylum Cyanobacteria, was found in abundance in the ML4-I sample.

Microbial lineages that are likely associated with anoxic conditions and are primarily found in sediment samples (MS4-I, MS3-I, MS2-I, ML1-I). Within the phylum Chloroflexi, the genera AB-539-J10, GIF9, and *Thermoflexus*, which are known to include fermentative bacteria, were abundantly present in both sediments and microbialites of Zone I. Other genera such as *Acetothermiia*, *Aminicenantes* (Acidobacteria), *Cryptanaerobacter*, *Pelotomaculum* (Firmicutes), GAL15, and *Thiobacillus* (Gammaproteobacteria) were significantly more abundant in the sediments compared to other groups. The only identified Archaea phylum, Bathyarchaeota, was exclusively found in the sediment samples.

## DISCUSSION

### Environmental Controls on Microbialite Formation

Streams and groundwater feeding the entire catchment area of the lake and the lake itself are consistently rich in Mg and low in Ca content with slightly alkaline pH and there is no significant seasonal fluctuation in water chemistry (Table 2). A cation composition of Mg > K > Si> Ca> Na is considered typical for the lake catchment area (groundwater, springs and streams) while Mg > Na > Si > Ca >K represents the lake water chemistry in decreasing order (Table 2). Moreover, there is no visible carbonate precipitation throughout the flow paths of the springs and streams. Additionally, no thin layer of carbonate (e.g., carbonate ice) formation at the air-water interface was observed where the spring waters emerge. The saturation indices (SI) calculated by PHREEQC for Mg and Ca-bearing minerals (Table 5) indicate that the lake water is supersaturated with respect to hydromagnesite in the wet (March) and the dry season (August). Furthermore, the calculated saturation indices revealed supersaturation with respect to other Ca/Mg-carbonate minerals, such as dolomite, aragonite, calcite, huntite, and magnesite in both surface and groundwater, in addition to the lake water. However, despite the supersaturation of these carbonate minerals, there is a lack of carbonate precipitation, which appears to be related to kinetic effects, and kinetic inhibition. It is well established that Mg acts as a kinetic inhibitor for carbonate precipitation (Berner 1975). A typical example of a kinetic effect is widely observed in marine environments for the preferential formation of aragonite over calcite due to high Mg/Ca ratios. In contrast to the spring and stream water, extensive precipitation of hydromagnesite with minor amounts of aragonite is only observed in the lake. As revealed by the current and previous studies, hydromagnesite is the dominant hydrated Mg carbonate mineral in both the actively growing and sub/fossil microbialites throughout the entire lake, regardless of the observed morphotypes and structural differences in the described zones (Figs.6,7 and 8). Palygorskite is another mineral identified in the lake, particularly in the thick mucilaginous orange-colored, organic rich microbial layer of the deeply growing thrombolite (at 15 m water depth) in Zone I and nesquehonite is only present in carbonate muds in the shoreline (Fig.3). Palygorskite is an Mg-rich phyllosilicate with an approximated formula of yMg_5_Si_8_O_20_(OH)2 · (1 − y)[xMg_2_Fe_2_ · (1 − x)Mg_2_Al_2_]Si_8_O_20_(OH)_2_. The presence of previously unidentified palygorskite mineral in the microbialites of Lake Salda could be attributed to detrital grains carried from nearby deltaic deposits and trapped into the microbial layer, or in-situ precipitation of Si released from the decomposition of Si- bearing EPS or dissolution of diatoms within the microbial layer (MDL1; Fig. 4D, Fig. 5F). The exact reason for the formation of palygorskite at the identified depth and zone in Lake Salda remains unclear and is worthy of further investigation. While aragonite formation is generally attribute to evaporative processes in lacustrine environments, its occurrence in Lake Salda at a water depth of 10 m, along with its association with the microbial layer collectively points to a microbially mediated origin. Based on the recent finding of Burnie *et al*. (2023) and Balci (2023), given its relative stability, aragonite would be a promising mineral for biosignature preservation in the Lake Salda microbialites.

**Table 5.**
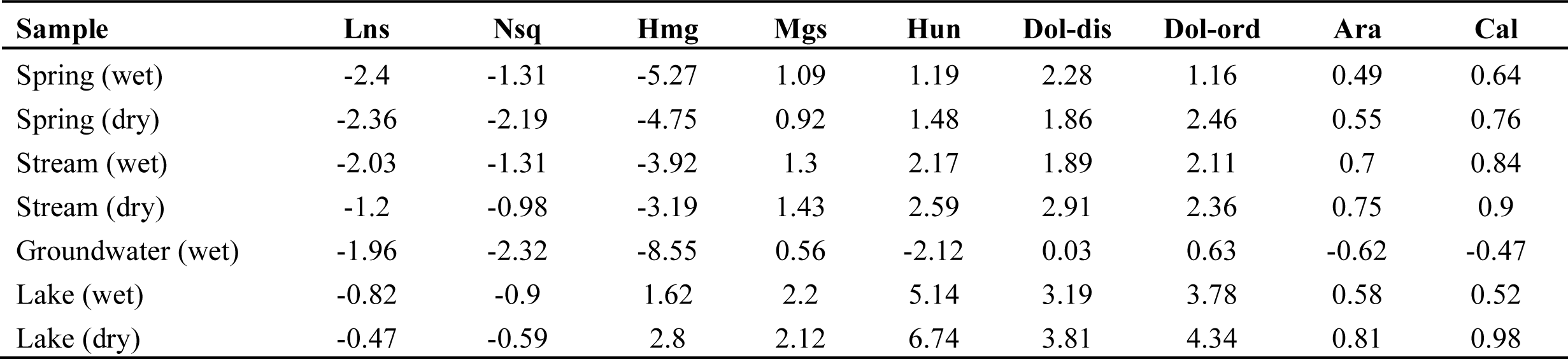
Saturation indices calculated by PHREEQC for relevant mineral phases including lansfordite (lns), nesquehonite (nsq), hydromagnesite (hmg), magnesite (mgs), huntite (hun), disordered dolomite (dol-dis), ordered dolomite (dol-ord), aragonite (ara) and calcite (cal).

It is widely accepted that environmental factors primarily control the overall microbialite morphology while the microorganism plays a key role in the accretion and fabric of carbonates (Riding 2000, Paul *et al*., 2016). In Lake Salda, both the actively growing and sub/fossil microbialite mounds are present within the extensive alluvial fan deltas (Salda and Doğanbaba) where the seepages of groundwater generally exist (Fig.1). The deltaic deposits, primarily composed of ultramafic rocks in various sizes (e.g., cobbles, pebbles) and ultramafic- derived sediments, likely promote microbialite growth by acting as both substrates as nucleation sites and source for cations such as Mg and Si. It is equally possible that the deltaic deposits and groundwater provide the key nutrients to the lake ecosystems since the lake is oligotrophic (P-N-limited) (Kazanci *et al*., 2004). Another key contribution of the alluvial fan deltas to microbialite development is to transform a deeper shorelines of the lake into a shallow and occasionally restricted embayment where microbialites started to accumulate. Given the lack of significant fluctuations in the water chemistry of the lake including pH, and cation compositions over the years, spatial differences observed in the morphotypes of the microbialites are likely developed as a response to changes in lake level, detrital input, and sedimentation rate. Additionally, high wind activity and wave erosion seen in the lake could possibly agitate the shorelines, thus keep the particles suspended in the shorelines.

A notable difference observed in the shallow versus the deeper grown microbialites in Zone I is a typical example of environmental controls on the microbialite morphology. For example, the columnar-linked thrombolites exclusively exits at 10-20 m water depths Zone I (Fig. 6C) while the dome-shaped thrombolites with cauliflower morphology or stacked dome-shaped thrombolites and stromatolitic-thrombolites are the common morphotypes for the shallow growing microbialites (Fig.6). In contrast to the shallow growing ones, the upper surface of the deeply growing thrombolites is covered by the well-developed commonly interconnected finger-shape mini columns (a few cm in size) (Fig. 6C,D; Fig 7E). These mini-columns represent visible internal layers Fig. 7E) as well as grainy fabrics with no layers (Fig. 4C)). It is reasonable to assume that these deeply growing microbialites (> 10 m) are less affected by the water level fluctuation (Kazanci *et al.,* 2004), evaporation and wind/wave agitation, enabling to development and protection of thick and proliferated microbial layers that produce organic matrix/EPS as a nucleation site for the precipitation (Fig. 2 and Fig. 7E). Extensive presence of stalk producing diatoms and occurrences of hydromagnesite crystals within the EPS are further support this hypothesis. Diatoms play a large role for the production of organic matrices that are primarily composed of sulfated polysaccharides, proteins, and some uronic acid over multiple centimeters thick (Somanader *et al*., 2022). The available sunlight reaching the deeper level of the lake can enable these photosynthetic species to colonize. Even though the same microbial layers including diatoms exist in the shallow microbialites, the lack of such structures on the near shore or shallow microbialites suggests that these structures were likely truncated by the wave and high wind activity in the lake. This seems an effective post or syn/depositional process since their debris renders much of the shoreline white.

Closer observation of the dome or half-domed shape microbialites revealed differences in laminations and appearances at their uppermost surface. At the meso scale, cauliflower top stromatolitic–thrombolites and calcareous-knobs-top thrombolites that are usually growing upwards are common in Zones I, II and IV (Fig. 6, Fig. 7A-C). In contrast, the stacked-domed shape microbialites with the smooth surface texture in Zone VI show well-developed radially growing stromatolitic layers (Fig. 8A-C).

### Microbial Diversity of Microbialites

Earlier studies of Lake Salda microbialites have simply documented the microbialites and their association with plausible microbial activity (e.g., diatoms and algae) (Zedef and Braithwaite, 1996). A few molecular studies have been performed to decipher the microbial diversity of the microbialites; which are primarily restricted to the near shore microbialites in the southwestern part of the lake and the sediments (Poyraz and Mutlu 2017; Balci *et al*., 2020; Balci 2023). The present study provides first comprehensive coverage of the microbial populations of the microbialites existed in both the different zones and the water depths (Table 1). With these analyses, we were able to generate a more comprehensive survey of the microbes present within the different microbialite types.

Genetic data revealed that the relative abundance of major taxa differs substantially between the zones and water depths in Lake Salda. For example, in Zone I, bacterial abundances vary between the shallow and the deeply growing thrombolitic microbialites as well as the lake sediments (Fig. 9). Notably, the thrombolitic microbialites at water depths greater than 5 m (ML3-I; ML4-I and ML5-I) show a high percentage of *Exiguobacterium* genus, which belongs to Firmicutes phylum (Figs. 9 and 10). Surprisingly, *Exiguobacterium* spp. are particularly enriched in the orange-colored mucilaginous layer of the microbial mat at a depth of 10 meters (MD-L1, Fig. 2D and Fig. 4C), comprising up to 93 % of the bacterial population.

Like Zone I, *Exiguobacterium* spp. is consistently present on the upper surface of all microbial mats with varying abundance (0.1- 93 %) throughout the entire lake, in particular, at meter’s depth (ML3-I, 5 m; ML4-I, 10 m, and ML5-I, 15 m). Together these data suggest that *Exiguobacterium* spp are a numerically significant component of the Lake Salda microbialites. Scanning electron microscopy (SEM) observations of microbial layer with high abundance of *Exiguobacterium* genera also showed a diverse range of diatoms (Fig. 4D, E). As in the current study, *Exiguobacterium* strains have been previously identified and isolated from various microbialite sites, such as cold freshwater thrombolites (Gutiérrez-Preciado *et al*., 2017), a warm (i.e., ∼24°C) freshwater stromatolite in Lake Socompa (Ordoñez *et al*., 2015) and Pavilion Lake, Canada (at 20 m water depth) (Whiteet *et al*., 2019). There are several proposed ways in which *Exiguobacterium* spp might contribute to microbialite formation and maintenance. One of them is to production of biofilm. In nutrient-poor oligotrophic conditions like Lake Salda, *Exiguobacterium* spp. are known to produce copious amounts of biofilms as a survival strategy (Gomez-Javier *et al*., 2018). These biofilms may act as structuring agents in microbialite formation by adhering to precipitated carbonates or providing nucleation sites for mineral deposition.

*Exiguobacterium* spp. also exhibit ureolytic activity, leading to an increase in water alkalinity and promoting carbonate precipitation (White *et al*., 2019). Although the alkaline characteristics of Lake Salda water may render this metabolic contribution to microbialite development insignificant, the high abundance of aragonite within the microbial layer (MLD- 1) of deeply growing microbialites (>10 m), where the entire bacterial community is dominated by *Exiguobacterium*, suggests the potential involvement of this metabolic activity in the lake. Genus of the *Exiguobacterium* has been shown to induce aragonite formation as a result of the degradation of Cyanobacteria and extracellular organic matrix (Pace *et al*., 2016). Furthermore, *Exiguobacterium* spp are known to synthesize various compounds such as carotenoids and orange pigments, which contribute to the visible color of the microbial layers in Lake Salda. These compounds provide protection against ultraviolet radiation, and *Exiguobacterium* spp. also produce indole-3-acetic acid (IAA), known to promote the growth of microalgae and diatoms (Amin *et al*., 2015; Lee *et al*., 2019). In the oligotrophic environment of Lake Salda, these beneficial relationships are likely significant, providing not only nutrient sources but also protection from UV radiation. Despite these insights, the exact role of *Exiguobacterium* spp. in the development of microbialites in Lake Salda remains elusive, requiring further investigation. Compared to the other zones, cyanobacteria identified in Zone I except for the sample ML4-I, regardless of the water depth, is extremely low (e.g., 0.2 %; Fig.9). However, it should be noted that the total percentage of cyanobacteria detected through microbial community analysis does not correlate with the volume that occupies in the microbial mats due to the relatively larger cyanobacteria cell. In contrast, the phylum Chloroflexi, were identified in all sample types (sediment and microbial mats) as one of the core populations, and for most of the cases, the dominating proportion of microbial community, in particular those of the stromatolitic, stromatolitic-thrombolites in Zones III, IV and VI (13-34 %) (Figs.9, 10). The Chloroflexi present in the Lake Salda mats fall into the class Anaerolineae and Dehalococcoidia with dominant genera of SBR1031 and A4b a group which is typically characterized by anaerobic, non-phototrophic heterotrophs (Yamada *et al*., 2006), though many appear to have ability for aerobic respiration (Hemp *et al*., 2015a,b; Pace *et al*., 2015; Ward *et al*., 2015a,b). Thus, we suggest that the Chloroflexi identified in the mats could play a key role in both aerobic and anaerobic organic matter mineralization, thus breaking down biomass produced by Cyanobacteria and other photoautotrophs. These classes of bacteria have been recently reported to be involved in carbonate precipitation via increasing alkalinity (Paul *et al*., 2016; Ward *et al*., 2020; Okumura *et al*., 2022).

Compared to Zone I, Cyanobacteria are more abundant in Zones III, IV, V and VI where stromatolitic thrombolites are more common and make up 14 -40 % of the microbial community (Figs.9 and 10). The dominant Cyanobacteria are members of the genera *Coleofasciculus*, *Leptolyngbya, and Nodosilinea*. Genus of filamentous *Phormidium* and genus *Leptolyngbya* are only abundant in Zones III, IV V and VI. Unlike the other genera of Cyanobacteria, genus *Coleofasciculus* is more abundant in the microbial layers of a thrombolite at a water depth of 15 m (MD-L2 and MD-L4) in Zone I (Fig. 2D and Fig. 10).

Similar to Lake Salda, Cyanobacteria genera of *Coleofasciculus*, *Leptolyngbya, and Nodosilinea* were reported in the microbialites in Japan (Shiraishi *et al*., 2017). Furthermore, as we observed here, a link between morphotypes of microbialites and cyanobacterial phylotype compositions was revealed. Accordingly, *Leptolyngbya* spp. were abundant in thrombolitic structures while *Phormidium* spp. were more associated with stromatolitic structures in the Ueno microbialite site in Japan. It is well documented that cyanobacteria play a major role in the development of microbialites by trapping and binding processes in addition to in-situ precipitation (Dupraz *et al*., 2009).

OTUs belonging to the gammaproteobacterial genus *Thiobacillus* that include, anoxygenic phototrophic sulfide-oxidizing bacteria, were only present at 0.6 -6.5% abundance in the microbial mat obtained from the shoreline sediments of Zone I and III where the groundwater seepages are present (Table 4). This suggests likely using the photosynthetic oxidation of sulfide produced by the class of Desulfobacterota -sulfate reducers- identified in the sediment (MS1-I; MS2-I and MS3-I) to drive carbon fixation. (Wilbanks *et al*., 2014). This type of anaerobic closed internal sulfur cycle in the lake only present in the shoreline sediments of Zone I and III. *Brevundimonas, Hyphomicrobium* and *Hydrogenophaga* (Proteobacteria), and *Luteolibacter* (Verrucomicrobiota), are the other genera found abundantly in Zone I, particularly in the shallow-grown microbialites.

### Microbial Effects on Hydromagnesite Precipitation

Among the observations at Lake Salda microbialites, the spatial distributions of heavily and poorly calcified filamentous cyanobacteria in addition to mucilaginous thick organic matrix/EPS in the upper surface of all microbialites seem to be critical toward understanding the microbial carbonate formation mechanism.

As indicated by the thermodynamic calculations from the water chemistry data, Lake Salda water is saturated with respect to various carbonate minerals including hydromagnesite, magnesite, dolomite and slightly saturated with aragonite and calcite (Table 5). Despite the supersaturation, mineralogical characterization of the microbialites as well as other carbonate phases in the lake showed that hydromagnesite is the dominant carbonate mineral with a minor amount of aragonite. Although the other carbonate phases are supersaturated, the lack of precipitation is presumably due to kinetic inhibition. It is well established that the initial nucleation of carbonate minerals often enforces the largest kinetic barrier to the precipitation of these mineral phases from supersaturated solutions. Organic molecules play a key role by drastically reducing the kinetic barrier and controlling the mineral precipitation phases.

Genetic data and microscopic observations of the microbial mats and layers in Lake Salda showed variable numbers of coccoid and filamentous bacteria (e.g, Cyanobacteria, Chloroflexi), diatoms, and heterotrophic bacteria embedded in an organic matrix/EPS in which hydrated Mg-carbonate precipitation takes place (Figs.4 and 5). The aggregate of authigenic carbonate grains, cyanobacterial filaments, and diatoms typically constitute a complex porous network within the EPS (Fig. 5A,B), and micro- and nanocrystalline hydromagnesite is closely related to the organic matrix and filamentous cyanobacteria (Fig. 5C,D). However, it appears that not all cyanobacterial filaments are encrusted, and some filaments are not calcified (Fig. 5E). Degree of cyanobacterial calcification can be primarily attributed to the difference in their exopolymer compositions (Shiraishi *et al*., 2017). For example, *Leptolyngbya* spp. which secrete non-acidic polymers is found to be poorly calcified, forming peloids, thus the clotted fabric of thrombolites, while *Phormidium* sp. secreting acidic exopolymers is heavily calcified, producing the laminated fabrics of stromatolites. This finding is suggestive that exopolymer properties of different cyanobacterial taxa have control on microbial carbonates fabrics. Such similar microbial fabric control is certainly possible in Lake Salda since Cyanobacterial genera secreting both acidic and non-acidic exopolymers were identified in the lake (Figs. 9 and 10). In fact, abundance at the mesoscale, stromatolitic thrombolites with the laminated fabric are present in Zones where microbial community is dominated by cyanobacteria (Fig. 8A, B and C).

In general, hydromagnesite crystals within the EPS show spherulitic growth fabric typically formed by the arrangement of the plate and blade-like crystals in a radial or non-oriented way (Fig. 5B-D). In accordance with the present study, spherulitic carbonates that grew in the presence of various organic compounds, such as carbohydrates and amino acids, as an analog for the effect of EPS in biofilms, are reported (Braissant *et al*., 2003). These hydromagnesite spherulites typically ranged from 5 and 20 µm in size and are often aggregated within the EPS-rich zone (Fig. 5D). Micro and nano/hydromagnesite crystals that encrust filamentous cyanobacteria present as plate and blade-like crystals (Fig. 5C,D). Small spheroidal carbonates with a size of <1 µm is also present as incrustation of the filaments. Energy dispersive X-ray analyses (EDS) revealed the presence of C, O, Mg, Ca, Si and S (Fig. 5F). Carbonates are also found as bigger clusters embedded in the EPS matrix (Fig. 5B).

The prevalence existence of carbonate particles/aggregates within organic/EPS-rich microbial mats of Lake Salda suggests that carbonate precipitation takes place under the influence of an EPS matrix, thus organo-mineralization- in which EPS likely acts as a substrate and controls carbonate nucleation and crystal growth which is kinetically inhibited in the lake. Similar to Lake Salda, in modern as well as ancient carbonate microbialites, irregular micrite laminae or peloids composed of micro/nanocrystalline, carbonates are considered to be produced by precipitation related to organic polymers and with strong biological influence (Riding 2011). Presence of hydromagnesite crystals within thick EPS and remnants of filaments within Lake Salda microbialites refer to the effect of organic matrix/EPS on precipitation from supersaturated solution, possibly influenced locally by microbial activity. Compared to the stromatolitic structures, the relatively abundant heterotrophic bacteria in the microbial layers of the thrombolitic structures in Zone I emphasize the role of organic matrix/EPS for carbonate fabric development of the microbialite in the lake. Degradation of EPS by the bacteria changes the chemical structure of the extracellular matrix, modify functional groups and the acidity of the surface, thus modify the organic template that precipitation occurs (Park *et al*., 2014; Karami *et al*., 2014). Such processes may readily prevent the formation of continues layers but give rise to patchy carbonate precipitation that we observed in Zone I (Fig. 7A). Thereby, characterizing and quantifying of organic matrix/EPS composition of each microbial layer is critical for the mechanistic understanding of the organo-mineralization processes (the influence of organic matrix, EPS, etc.) in the lake.

## CONCLUSION

In Lake Salda, despite a common external dome-shaped or stacked dome-shaped cauliflower macromorphology, thrombolitic, stromatolitic-thrombolites and to a lesser extent dendritic mesoscale internal growth structures are identified at separate locations and water-depth within the lake.

The current study involved comprehensive molecular analysis of microbial mats utilizing Illumina sequencing, which successfully identified at least 12 dominant bacterial phyla. Thrombolitic, in particular those in the deeper part (< 5m water depth) and stromatolitic structures present in the lake are distinguished by their microbial communities. Of particular significance, Anaerolineae (the genera A4b and SBR1031 genus) and Cyanobacteria (the genera *Cyanobacteriales, Leptolyngbyales, Phormidesmiales*, and *Synechococcales*) emerged as core populations, prominently present in the microbialites with the stromatolitic and stromatolitic-thrombolitic growth structures that are present in the shallow waters (< 5 m) (Zones III, IV, and VI). Conversely, within the phylum Firmicutes, *Exiguobacterales* order and genus *Exiguobacterium spp.* consistently dominated the thrombolitic microbialites, representing a substantial proportion of the microbial communities of the deeper-growing ones (MDL, ML3-I, ML4-I, ML5-I). The noteworthy prevalence of *Exiguobacterium spp*. solely within the deeply growing thrombolites can be attributed to the presence of abundant organic matter, which was likely shielded from UV decomposition and water agitation.

Overall, a commonality among microbialites is the presence of the phyla Chloroflexi, Firmicutes, Cyanobacteria and Proteobacteria. This microbial community profile of microbial mats and concurrent presence of diatom-rich layers seem to play a pivotal role for the construction of microbialites by governing the cycling (production vs. degradation) of EPS, in which carbonate precipitation abundantly takes place.

Existence of fossil/sub-fossil and modern microbialites within the alluvial fan deposits, predominantly composed of ultramafic rocks of various sizes, indicates the availability of substantial substrates as nucleation sites. Furthermore, physicochemical factors, such as increased cation release (e.g. Mg and Si) arising from enhanced water-rock interaction have significantly promoted the growth of microbialite in the lake.

Microscopic analysis of the microbial mat unveiled a notable association of nano/microcrystalline platy and sub/spherical hydromagnesite crystals with the organic matrix/EPS and filamentous cyanobacteria sheaths. The frequently identified spherical crystals within EPSs can be explained by microbial mediation that becomes the controlling factor of hydromagnesite precipitation only when precipitation is kinetically inhibited. We propose that the macro morphology of the microbialites is largely controlled by environmental factors such as lake-level while mesoscale growth structure is primarily controlled by the microbial community composition of the mats.

